# Loss of Sarm1 non-autonomously protects Schwann cells from chemotoxicity

**DOI:** 10.1101/493163

**Authors:** Weili Tian, Tim Czopka, Hernán López-Schier

**Affiliations:** Sensory Biology & Organogenesis, Helmholtz Zentrum Munich, Germany; Institute of Neuronal Cell Biology, Technical University of Munich, Germany

**Keywords:** Wallerian degeneration, Sarm1, Schwann cells, regeneration

## Abstract

The obligate pro-degenerative protein Sarm1 is essential for Wallerian axon degeneration. Inhibition of Sarm1 has been proposed as a promising neuroprotective strategy with clinical relevance. Yet, the conditions that will most benefit from inhibiting Sarm1 remain undefined. Here we use genetics and pharmacology in zebrafish to show that systemic elimination of Sarm1 is glioprotective. Loss of Sarm1 does not affect macrophage recruitment to the wound microenvironment, focal injury resolution, or nerve repair. Unexpectedly, Sarm1 deficiency increases Schwann-cell resistance to toxicity by diverse chemotherapeutic agents after neuronal injury. Yet, synthetic degradation of Sarm1-deficient severed axons reversed this effect, suggesting that glioprotection is non-cell-autonomous. These findings anticipate that interventions aimed at inhibiting Sarm1 can counter heightened glial vulnerability to chemical stressors and may be an effective strategy to reduce chronic consequences of neurotrauma.

## INTRODUCTION

The peripheral nerves that communicate skin, muscle and sensory organs with the brain must maintain functionality throughout life despite frequent stress and trauma (1–6). Loss of integrity of peripheral neurons and associated cells, including glia, is a common occurrence in severe neurological dysfunctions that include weakness, pain and loss of sensation (7). Glial loss leads to nerve demyelination, defasciculation and neuronal death. Nerve injury triggers axon fragmentation and degeneration. In turn, this induces the dedifferentiation of associated glial cells, which enhances glial role in nerve repair. However, dedifferentiation also makes glia vulnerable to degeneration after protracted denervation (8–12). Damaged axons undergo Wallerian degeneration (13), which is regulated by the evolutionary ancient pro-degenerative protein Sarm1 (<u>S</u>terile <u>A</u>lpha and TI<u>R</u> <u>M</u>otif containing 1) (14–17). Sarm1 is a modular protein that contains two sterile-alpha motifs (SAM), one mitochondrial association (MT) and one TIR domain (14). Nerve injury activates Sarm1 by TIR dimerization, which is sufficient to induce the axon degeneration (18, 19). Mechanistically, Sarm1 activation results in the loss of nicotinamide adenine dinucleotide (NAD+) in damaged axons (20–22). The TIR domain acts as a NADase (23), explaining why over-expression of the NAD+ synthesizing enzyme NMNAT1 inhibits the degradation of severed axons by sustaining high levels of NAD+ downstream of Sarm1 (24). Moreover, the loss of Sarm1 blocks the degeneration of injured axons, and forced activation of Sarm1 induces axon destruction in the absence of injury. Therefore, Sarm1 is a cell-autonomous hierarchical regulator of a signaling pathway that is necessary and sufficient for axonal degradation (22). Upon peripheral-nerve injury, glial Schwann cells acquire a specialized function that promotes the clearance of axon fragments ahead of axonal re-growth from the proximal stump that remains associated to the neuronal perikaryon (25). Expeditious axonal regeneration is important because sustained Schwann-cell denervation leads to protracted loss of glial terminal phenotype and eventual death. In turn, glial loss impairs regenerating nerve myelination and circuit repair, transforming acute neuropathies into irreversible chronic neurological dysfunction. Therefore, inhibiting or delaying axon destruction has been hypothesized as an effective strategy to counter heightened Schwann-cell vulnerability to additional stressors that include metabolic imbalance and drugs (26, 27). This idea has sparked intense efforts to identify specific molecular inhibitors of axon degeneration for clinical applications (28, 29). As a discrete hierarchical factor that is essential for axon degeneration, Sarm1 is an ideal target for pharmacological interventions (18). However, the conditions that would most benefit from inhibiting Sarm1 systemically are yet to be defined (30, 31).

Here we address the above issue using the powerful genetics of zebrafish, a small vertebrate whose nervous system is anatomically simpler but structurally and functionally similar to that of mammals (32–34). Crucially, the zebrafish larva is ideal to study Schwann-cell biology in the natural context of the behaving animal (35–42). By characterizing loss-of-function mutations in Sarm1, we provide novel insights into the cellular basis of axon-glia interactions, and provide data that encourage the quest to identify and develop Sarm1 inhibitors for clinical applications.

## RESULTS

### Identification and mutagenesis of Sarm1 in zebrafish

The amino-acid sequence of Sarm1 is well conserved across species (15, 43). To identify Sarm1 orthologs in zebrafish, we scanned publicly accessible genomic data (*Danio rerio* reference genome assembly version GRCz11) by a BLAST search using the TIR domain, which is present in all known Sarm1 proteins (15, 44). This exploration yielded a single candidate *locus* in chromosome 15. No other part of the zebrafish genome appears to harbor Sarm1 paralogs. The genomic structure of the putative zebrafish Sarm1 reminisces that of other species, containing 8 exons that code for a protein of 713 amino acids, with the typical N-terminal auto-inhibitory domain, two central SAM multimerization domains and a C-terminal TIR degeneration domain (Figure 1A). Similar to *D. melanogaster*, however, *D. rerio* Sarm1 lacks an obvious mitochondria-targeting sequence (MT). To test whether the identified gene produces a protein with the expected functional role, we used CRISPR/Cas9-mediated genome modification to generate loss-of-function mutations in Sarm1. By targeting exon 1, we obtained germ-line transmission of two alleles: *sarm1^hzm13^* and *sarm1^hzm14^* (Supplemental Figure 1A-B). The hzm13 allele introduces an 11-base deletion and T/C mutation, resulting in a frame shift and premature stop codon. hzm14 is a 7-base deletion and AG/GA mutation that also generates a frame shift and premature stop codon. Analysis of protein extracts from wild-type embryos by Western-blot using an antibody to Sarm1 revealed a single band of approximately 80kDa, which agrees with the expected size of the full-length protein (Supplemental Figure 1C). This band was absent in protein extracts from homozygous *sarm1^hzm13^* zebrafish embryos. Of note, because this antibody recognizes an epitope in the C-terminus of Sarm1, it does not allow to discriminate between the expression of a truncated protein lacking all the domains with known function, and the complete absence of Sarm1 induced by nonsense-mediated mRNA decay. Homozygous *sarm1^hzm13^* mutants display no overt anatomical defects (Supplemental Figure 1D-E), are viable and develop into fertile adults (not shown). Furthermore, a simple assay for sensorimotor function that consists of eliciting the escape response after tactile stimuli showed that the displacement distance and the average acceleration were no different between wild type and Sarm1 mutants (Supplemental Figure 1F-G) (45).

**Figure 1.**
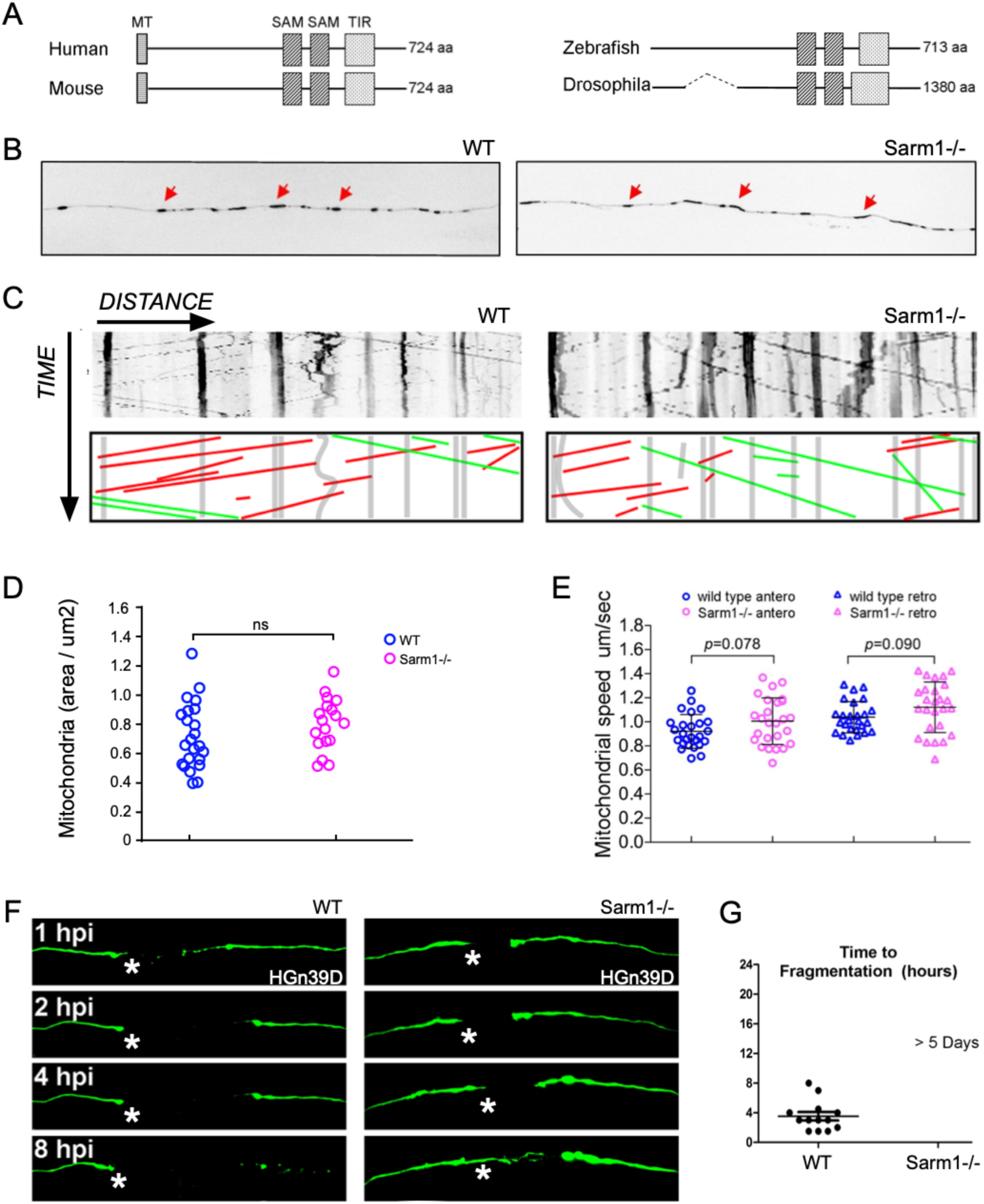
**A)** Structure Sarm1 indicating alignment of the Sarm1 functional domains from different species (not at scale). **B)** Confocal image of axonal mitochondria marked with mito-mCherry in wild-type and Sarm1-/-. Red arrows point to prominent mitochondrial groups in axons. **C)** Upper panels, kymographs from videomicroscopic recording of axonal mitochondria in wild type (H) (left panel) and Sarm1-/- (I) (right panel). Lower panels show color-coded traces of moving mitochondria in anterograde (green) and retrograde (red) directions, taken from the kymographs shown in the upper panels. **D)** Density of mitochondria in 5dpf wild type and Sarm1-/-, Error bar = SEM; n.s. = not significant, p value from Student’s t test, n=25 (WT), n=19 (Sarm1-/-). **E)** Mobility of the mitochondria in 5dpf wild type and Sarm1-/-. Circles show the anterograde and triangles the retrograde movement of the mitochondria. p value from one-way ANOVA, wild type n=26, Sarm1-/- n=26. **F)** Time-lapse images of axonal degeneration of GFP-labeled lateralis sensory neuron in wild-type (left) and Sarm1-/- larvae (right). hpi = hour post injury, scale bar=50μm, white asterisk indicates the regrowing axons from the proximal stump. **G)** Quantification of the time from axon transection to fragmentation in wild type (n=13) and Sarm1-/- (n=13).

### Functional conservation of Sarm1 in Wallerian axon degeneration

Although Sarm1 has already been extensively studied in neurons of *Drosophila* and mammals, we wanted to assess the effects of systemic loss of Sarm1 on neuronal and non-neuronal cells in larval zebrafish. The reason is that no single study has addressed Sarm1 function holistically, in a single organism, by *in toto* live microscopic imaging. To this end, we used several parameters of nervous-system structure and function by combining *sarm1^hzm13^* with various transgenes expressing fluorescent markers in the sensory neurons of the mechanosensory lateral line, as well as in their associated Schwann cells (46, 47). The lateral-line system is ideal for *in vivo* studies at high resolution over extended periods, and for physiological studies under normal and altered conditions (48–51). It combines the organization of a typical vertebrate sensory system with the amenability for controlled experimental interventions that include microsurgery, pharmacology and optogenetics (41, 42, 52, 53). We found that loss of Sarm1 does not affect the development and maintenance of the lateral-line sensory pathway (Supplemental Figure 1H-N). Next, we assessed neuronal intracellular dynamics and physiological function. To this end, we fluorescently marked mitochondria in lateralis neurons by expressing the mitochondria targeting sequence of the Cytochrome-C oxidase subunit 8A fused to mCherry (54). We chose mitochondria because mitochondria produce energy for the neurons and also regulate calcium levels, which are critical for the function and viability of axons and neurons. These intracellular organelles are dynamic and distribute throughout the neuron by active transport mediated by molecular motors that move along microtubule tracks. In axons, microtubules are polarized such that their plus ends are directed toward the axon terminals, in turn orienting the movement of distinct molecular motors in the antero- and retro-grade directions (55). Intra-axonal movement direction also reflects mitochondrial fitness because stressed organelles are biased in the retrograde direction (56, 57). Therefore, axonal mitochondria represent an optimal proxy for neuronal polarization, intracellular dynamics and overall cellular health. We found a qualitatively similar number, density and distribution of mitochondria in the peripheral axons of wild type and Sarm1 mutants. Kymographic analysis revealed a majority of static large mitochondrial groups, and some smaller fragments moving persistently substantial distances at constant velocity in the anterograde and retrograde directions (Figure 1B-C). Importantly, quantifications showed no significant differences in the number and spatial distribution of axonal mitochondria (Figure 1D), or movement velocity and direction between wild-type and mutant animals (Figure 1E).

Next, we assayed functional conservation of Sarm1 in this neuronal pathway using a previously established bioassay of neurotrauma, which employs intact larval zebrafish, laser-mediated severing individualized axons by single-neuron fluorescent-protein expression, and high-resolution intravital microscopy (41, 58). Axotomy was done by focusing an ultraviolet laser beam to a discrete region of peripheral nerve. Upon severing, the distal axon segment quickly degenerates in wild-type specimens, whereas the proximal segment that remains associated with the neuronal perikaryon remains viable (Figure 1F). Kinetic analysis shows that severed Sarm1-deficient axon segments resisted degeneration for over 5 days. The degeneration-resistant phenotype was rescued by the transgenic introduction of a fluorescent-tagged full-length Sarm1 in mutant neurons (Figure 1G).

### Loss of Sarm1 does not affect focal damage resolution

Physical injury often results in inflammatory responses that recruit immune cells to the wound. After nerve injury, Schwann cells release chemoattractants for macrophages before axon fragmentation is evident. Injury-mediated production of reactive oxygen species (ROS) recruits inflammatory cells, including macrophages. Studies *ex vivo* using mammalian cultured neurons showed that Sarm1 acts downstream of mitochondrial ROS generation. Therefore, we decided to test *in vivo* if the loss of Sarm1 as well as the absence of Schwann cells would affect macrophage recruitment to the site of injury. To this end, we generated a transgenic line expressing membrane-targeted EGFP under the control of the macrophage-specific promoter Mfap4. We combined this line with Tg[SILL:mCherry] and mutations in Sarm1 or Erbb2. Loss of Erbb2 in zebrafish impairs Schwann-cell migration along lateral-line axons and leads to nerve unmyelination and defasciculation (53, 59). We first injured nerves with the laser and assessed macrophage behavior at high resolution. We found that the onset of recruitment and number of macrophages at the wound did not differ between wild-type specimens and Sarm1 or Erbb2 mutants (Figure 2A and Supplemental Movies 1-3). Macrophages arrived from various locations moved along the proximal and distal part of the axons in wild-type and mutant specimens. The retention time of macrophages at the proximal side of the wound was unaltered in Sarm1 or Erbb2 mutants, although on average slightly lengthened in the absence of Schwann cells (Figure 2B). In wild type, and Sarm1 and Erbb2 mutants, macrophages engulfed debris locally at the injury site, and did not appear to be involved in the degradation of the distal part of the severed axons (Supplemental Movies 1-3). We found, however, a significant increase in the size of the engulfed debris by macrophages in Erbb2-mutant animals (Figure 2C). These results reveal that the loss of Sarm1 does not affect focal damage resolution by macrophages, which are recruited to the wound independently of Schwann cells. We did notice discrete and highly mobile axon fragments that were not associated to EGFP(+) cells in wild-type and Erbb2-mutant animals, but not in Sarm1-deficient fish. This observation suggests that activated phagocytic cells other than macrophages and Schwann cells engulf and clear distal axon fragments after damage, and that this does not occur in Sarm1-/- animals.

**Figure 2.**
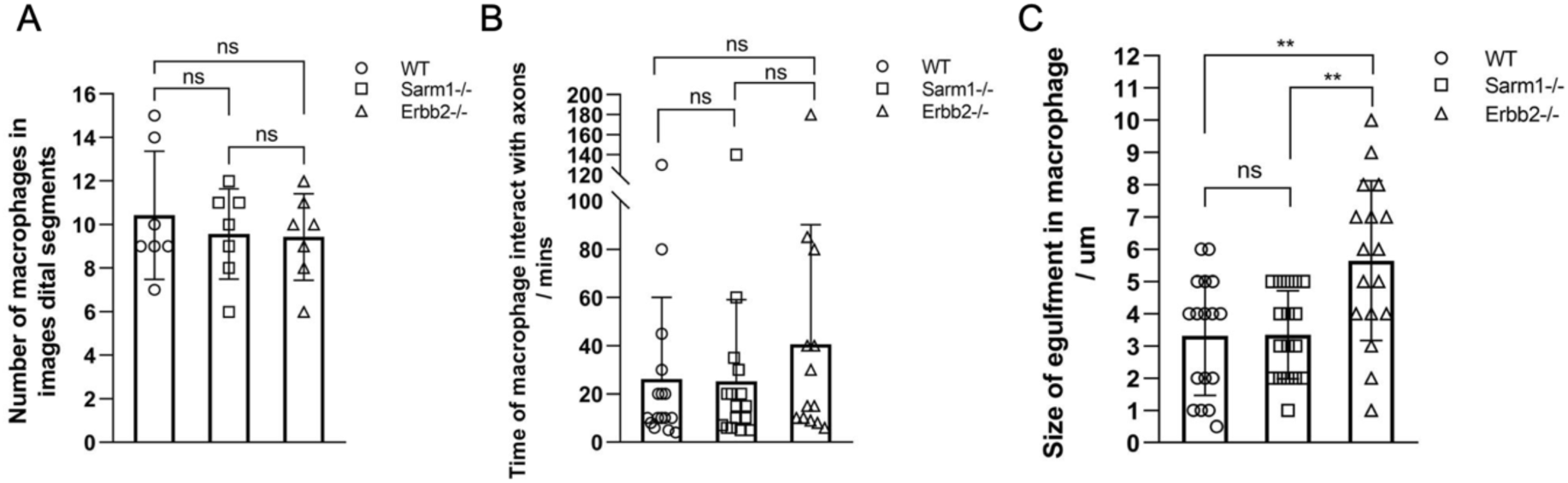
**A)** Quantification of the number of macrophages recruited to the injury site and adjacent axon segments. **B)** Quantification of the time macrophages interact with axon segments. **C)** Quantification of the size of debris within macrophages. P values for T-test.

### Synthetic elevation of axoplasmic Ca^2+^ induce degradation of severed Sarm1-deficient axons

Calcium (Ca^2+^) regulation in neurons is critical for homeostasis because sustained elevation of cytosolic calcium leads to axonal and neuronal degeneration. Often, this occurs because mitochondria release apoptosis-inducing factors and proteases in a calcium-dependent manner (60). Studies of mammalian neurons *in vitro* and of zebrafish have shown that neuronal damage triggers two waves of elevation of axoplasmic calcium (Ca^2+^) (21, 52). The first occurs nearly immediately after injury and decays rapidly, whereas the second has slower onset and decay. Loss of Sarm1 prevents Ca^2+^ elevation in severed axons (21). We sought to further test the functional conservation of Sarm1 in zebrafish *in vivo* by monitoring Ca^2+^ dynamics in lateralis sensory axons at high-resolution before and after injury. We measured Ca^2+^ dynamics *in toto* at high-resolution before and after axon injury using the genetically-encoded sensor GCaMP7a, to find that intact wild-type and Sarm1-mutant axons show undetectable levels of axoplasmic fluorescence above background (Figure 3A). Upon severing, fluorescent signal in wild-type axons distal segments increased immediately and subsequently decayed with a near constant slope, whereas fluorescence remained nearly undetectable in Sarm1-deficient distal axon segments (Figure 3B). Next, we examined the Ca^2+^ levels in mitochondria and the endoplasmic reticulum (ER) using genetically encoded vital sensors, respectively Mito-RGECO and ER-GCaMP3 (Figure 3C-F). We selected these organelles because Ca^2+^ release from the axonal ER activates the mitochondrial permeability transition pore to trigger axonal degeneration (61, 62). We found that both mitochondrial and reticular Ca^2+^ levels increased equally after axon severing in severed wild-type and Sarm1-/- axons. These results reveal that loss of Sarm1 attenuates calcium influx to the axoplasm, but not Ca^2+^ uptake in mitochondria or the ER.

**Figure 3.**
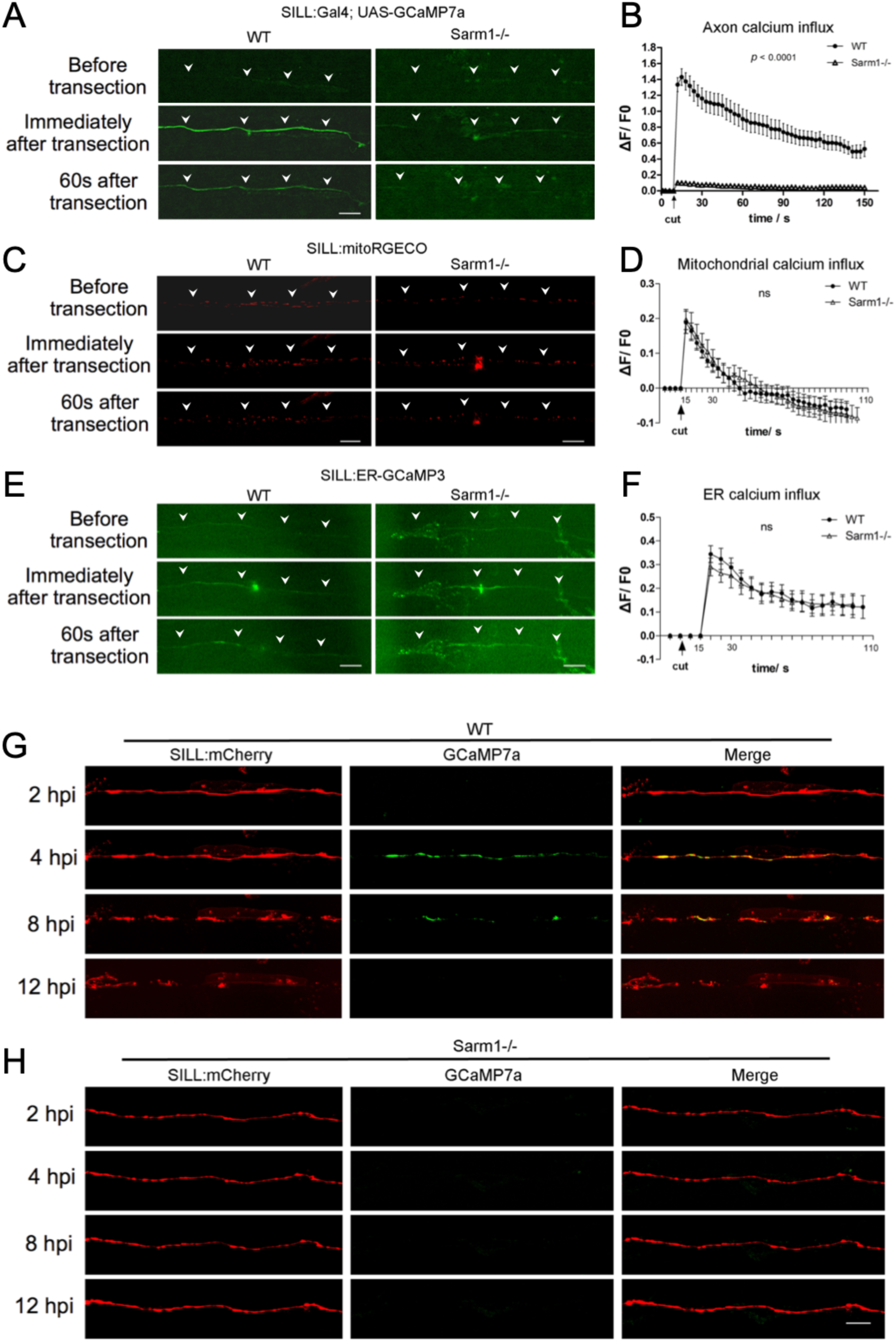
**A)** Confocal image of a single lateralis sensory axon expressing the green-fluorescent calcium sensor GCaMP7a in wild type (left column) and Sarm1-/- fish (right column). Rows show that same samples before laser-mediated transection (top), immediately after transection (middle) and 60 seconds after transection (bottom). In A, C and E, white arrowheads indicate the position of the axon, specifically when signal-to-background is low. Scale bar 20μm. **B)** Shows quantification of the first wave of axoplasmic calcium. Data are shown as mean ± SEM; p from one-way ANOVA, wild type n = 16, Sarm1-/- 16. **C)** Shows a confocal image of lateralis sensory axons expressing the red-fluorescent calcium sensor RGECO in wild type (left column) and Sarm1-/- fish (right column). Rows show that same samples before laser-mediated transection (top), immediately after transection (middle), and 60 seconds after transection (bottom). **D)** Quantification mitochondrial calcium influx shows the strong and nearly identical elevation and decay in wild type and Sarm1-/- immediately after the cuts. Data are shown as mean ± SEM; p from one-way ANOVA, wild type n = 16, Sarm1-/- 16. **E)** Shows a confocal image of lateralis sensory axons expressing the green-fluorescent calcium sensor CCaMP3 targeted to the endoplasmic reticulum (ER) in wild type (left column) and Sarm1-/- fish (right column). Rows show that same samples before laser-mediated transection (top), immediately after transection (middle), and 60 seconds after transection (bottom). **F)** Quantification ER calcium influx shows strong and statistically equal elevation and decay in wild type and Sarm1-/- after the cuts. Data are shown as mean ± SEM; p from one-way ANOVA, wild type n = 16, Sarm1-/- 16. **G)** Confocal image of a lateralis sensory axons expressing the mCherry (red) and green-fluorescent calcium sensor GCaMP7a (green) in wild type fish. Rows show same axons 2, 4, 8 and 12 hours after transection (hours-post-injury = hpi). **H)** Confocal image of lateralis sensory axons expressing the mCherry (red) and green-fluorescent calcium sensor GCaMP7a (green) in Sarm1-/- fish. Scale bar 20μm.

Because the second Ca^2+^ wave is responsible for the activation of the serine-threonine protease Calpain, which in turn facilitates axonal fragmentation by cleaving microtubules and neurofilaments (63), we decided to monitor Ca^2+^ levels in lateralis sensory axons 2, 4, 8 and 12 hours-post-injury (hpi). In severed wild-type axons, the second wave of axoplasmic Ca^2+^ starts 4 hpi in coincidence with axon fragmentation, and remains elevated in axonal debris up until 8 hpi (Figure 3G). Note that the temporal resolution of these images does not allow the resolution of degeneration before axonal regeneration. By contrast, the second wave of axoplasmic Ca^2+^ does not occur in Sarm1-mutant axons (Figure 3H). Thus, we hypothesized that forcing a sustained elevation of axoplasmic Ca^2+^ will be sufficient to trigger the degradation of severed Sarm1-deficient axons. To test this prediction, we transgenically expressed the rat transient receptor potential cation channel subfamily V member 1 (TRPV1) fused to tagRFP in lateralis afferent neurons of Sarm1-mutant zebrafish. Expression of mCherry alone in neurons served as control. TRPV1 is non-selective cation channel that exhibits a high divalent selectivity, and whose activation produces an influx of Ca^2+^ into cells (64). Rat TRPV1 is activated by temperatures above 43°C or by the vanilloid capsaicin. Importantly, this TRPV1 is inactive at the temperature used to maintain zebrafish (28°C), and zebrafish TRPV1 orthologs are insensitive to capsaicin (65). Therefore, rat TRPV1 expressed in zebrafish offers a tunable tool to elevate axoplasmic Ca^2+^ with excellent temporal resolution. We severed TRPV1-expressing and mCherry-expressing lateralis axons, and 2 hours later a vehicle solution or vehicle + capsaicin were added in the water holding the fish (Supplemental Figure 2A-B). Samples were inspected 90 minutes later. Severed Sarm1-deficient axons not expressing TRPV1 did not fragment in presence of capsaicin, or TRPV1-expressing axons bathed in ethanol solution. As hypothesized, we found that Sarm1-deficient TRPV1-expressing axon segments readily degraded in the presence of capsaicin (Supplemental Figure 2C-F). Thus, elevation of axoplasmic Ca^2+^ downstream of Sarm1 is sufficient to trigger axon degradation *in vivo*.

### Schwann cells are not essential to maintain Sarm1-/- axon segments

The Schwann cells support, fasciculate and myelinate sensory axons in vertebrates (66). We reasoned that delayed axon degeneration by loss of Sarm1 might impact the interaction between axons and Schwann cells. To assess this interaction *in vivo*, we combined the *sarm1^hzm13^* allele with the triple transgenic line *Tg[gSAGFF202A; UAS*-*GFP; SILL:mCherry]* to highlight the Schwann cells with green fluorescence and the lateralis afferent neurons with red fluorescence (53, 67). Using high-resolution intravital microscopy, we ascertained that Schwann cells develop normally and fasciculate sensory axons in Sarm1-deficient zebrafish (Figure 4A). Upon severing, wild-type axons were quickly cleared by the Schwann cells through engulfment of axonal fragments and intracellular degradation of debris (Figure 4B). However, there was no axon fragmentation or phagocytic activity by the Schwann cells in Sarm1 mutants (Figure 4C). Next, we asked if Schwann cells are necessary for the maintenance of severed Sarm1-deficient axons by generating a double mutant zebrafish line concurrently deficient for Erbb2 and Sarm1. In Erbb2-deficient specimens, the distal portion of the severed axons fragmented and were cleared (Figure 4C) but with a significant delay compared to wild-type specimens (Figure 4D). By contrast, in Erbb2/Sarm1 double mutants, severed axons did not fragment or degrade, identically to fish lacking only Sarm1 (Figure 4D). Thus, axon maintenance in Sarm1 mutants occurs independently of the Schwann cells, and that their eventual clearance is performed by other cells.

**Figure 4.**
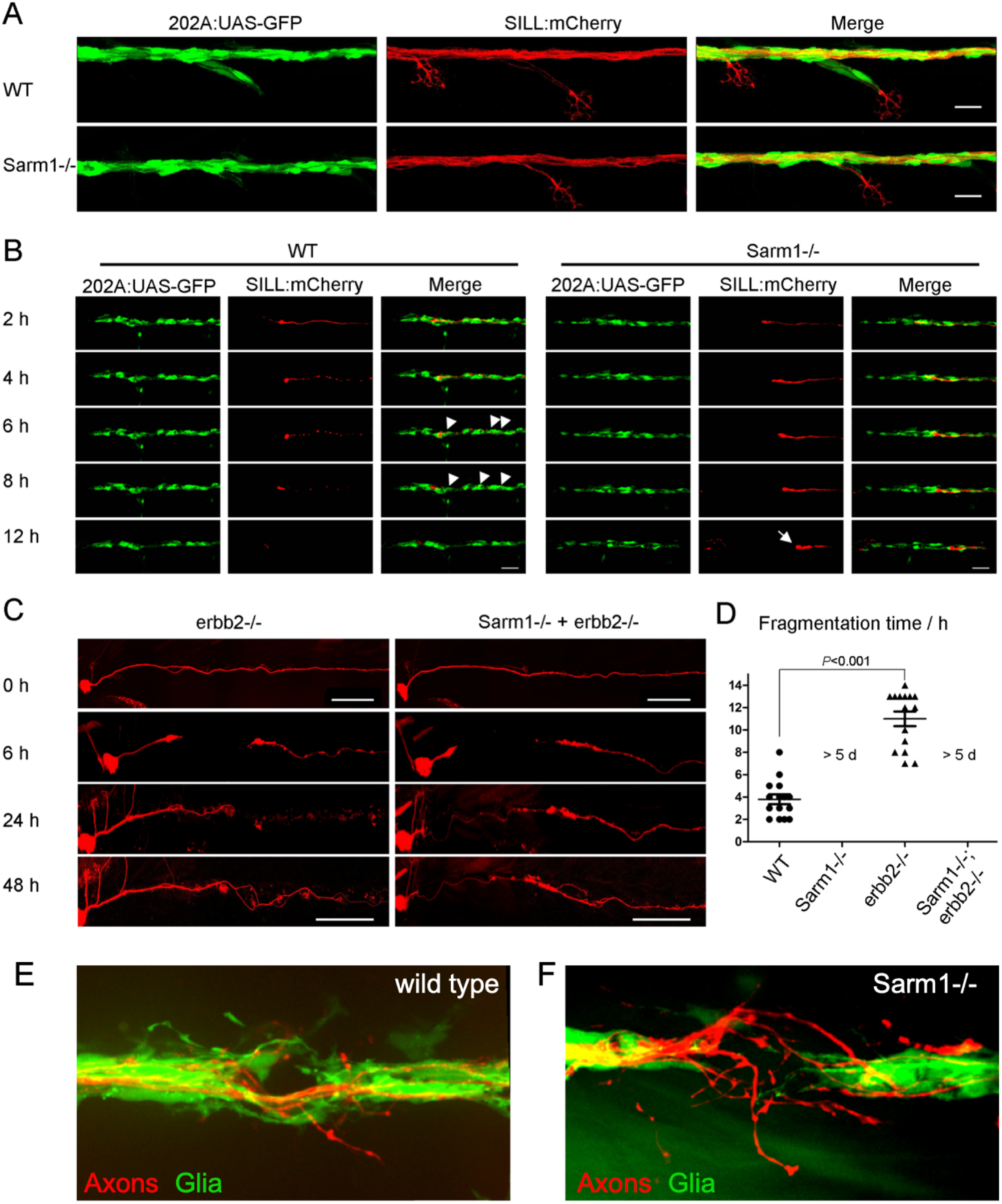
**A)** Confocal images of a double transgenic 5dpf larva showing Schwann cells marked by expression of GFP (green) under the control of the Tg[gSAGFF202A] Gal4 driver, and lateralis afferent neurons marked by expression of mCherry under the control of the SILL enhancer (red). Wild type (top), Sarm1 mutants (bottom). Scale bar 20μm. **B)** Images show the indicated time points after axon transection (hours post injury = hpi) from a videomicroscopic recording of Schwann cells (green) and their interaction with axons (red) in wild-type and Sarm1-/-. White arrowheads indicate Schwann cells engulfing axonal debris in the wild type. A white arrow indicates degradation-resistant axon segment in Sarm1-/-. Please, note that the proximal axon stump in Sarm1-/- is not visible in these images because it is outside the focal plane. **C)** Images of mCherry-expressing (red) transected axons in Erbb2-/- mutants and Sarm1-/-; ErBb2-/- double mutants. Scale bar 100μm. **D)** Quantification of transected axon fragmentation in Erbb2-/- and Sarm1-/-; ErBb2-/-. Error bar = SEM, p value from One-way ANOVA test, n = 15 (each group). **E)** Image of a from Supplemental Movie 1, showing the discrete local defasciculation of regenerated the sensory fiber (red) and the bridging of the glial gap by Schwann cells (green) in a wild-type specimen. **F)** Equivalent experiment, taken from Supplemental Movie 2, showing a more pronounced local defasciculation of the regenerated the sensory fiber (red) in a Sarm1-mutant specimen. Note that and the bridging of the glial gap does not occur.

When an injury generates a gap in the glia, wound-adjacent Schwann cells actively move and extend cellular projections resembling filopodia to quickly reconstitute a continuous glial scaffold (53). Although continuous glia are not necessary for the re-growth of the proximal axon stump, it prevents regenerating growth cones from straying, in turn facilitating end-organ *de novo* innervation and circuit reconstitution (39, 41). Thus, we decided to interrogate Schwann-cell behavior immediately after axon severing and during axon regeneration combining *Tg[gSAGFF202A;UAS-GFP; SILL:mCherry]* double transgene with the Sarm1 mutation. As expected, we found that in wild-type animals, the Schwann cells adjacent to the wound quickly extended filopodia to close the gap in the glial scaffold ahead of axonal regeneration (Supplemental Movie 4). Re-growing fibers then followed these sub-cellular glial bridges to reconstitute the nerve, suffering mild defasciculation restricted to a small area within the injury (Figure 4E and Supplemental Figure 3). In stark contrast, injury-adjacent Schwann cells did not migrate or produce filopodia-like projections in Sarm1 mutants (Supplemental Movie 5). Nonetheless, regenerating axons eventually negotiated the persistent larger gap to grow along the distal glial scaffold. However, the reforming nerves presented more pronounced local defasciculation (Figure 4F and Supplemental Figure 4).

### Loss of Sarm1 does not alter Schwann-cell phenotype after axotomy

Denervated Schwann cells undergo partial dedifferentiation from myelinating to a progenitor-like state as revealed by loss of expression of terminal-phenotype markers, including myelin and myelin-associated proteins (25, 68, 69). The loss of terminal phenotype promotes Schwann-cell proliferation and migration, which enhances their regenerative function (70). We hypothesized that because Schwann-cell denervation does not occur in Sarm1-mutants, distal Schwann cells may not dedifferentiate. This would explain their lack of phagocytic and protrusive activities after axon transection. Following this rationale, we immunostained samples with an antibody to Claudin-k, which localizes to the junctions between mature Schwann cells and is downregulated in denervated glia (41, 71). In wild-type specimens, Claudin-k remained strongly expressed along the entire length of the lateralis afferent nerve up to 6 hours after nerve injury (hpi), suggesting that distal Schwann cells remain mature during distal-axon fragmentation (Figure 5A-B). Beginning at 10 hpi, however, distal Schwann cells had markedly less Claudin-k than proximal cells. Finally, 24 hpi Claudin-k was conspicuously absent from distal Schwann cells even after axons regeneration had commenced. By contrast, Claudin-k remained strongly expressed after axon severing in Sarm1-mutant animals during the same period, with no apparent difference between Schwann cells located at either side of the wound (Figure 5A,C).

**Figure 5.**
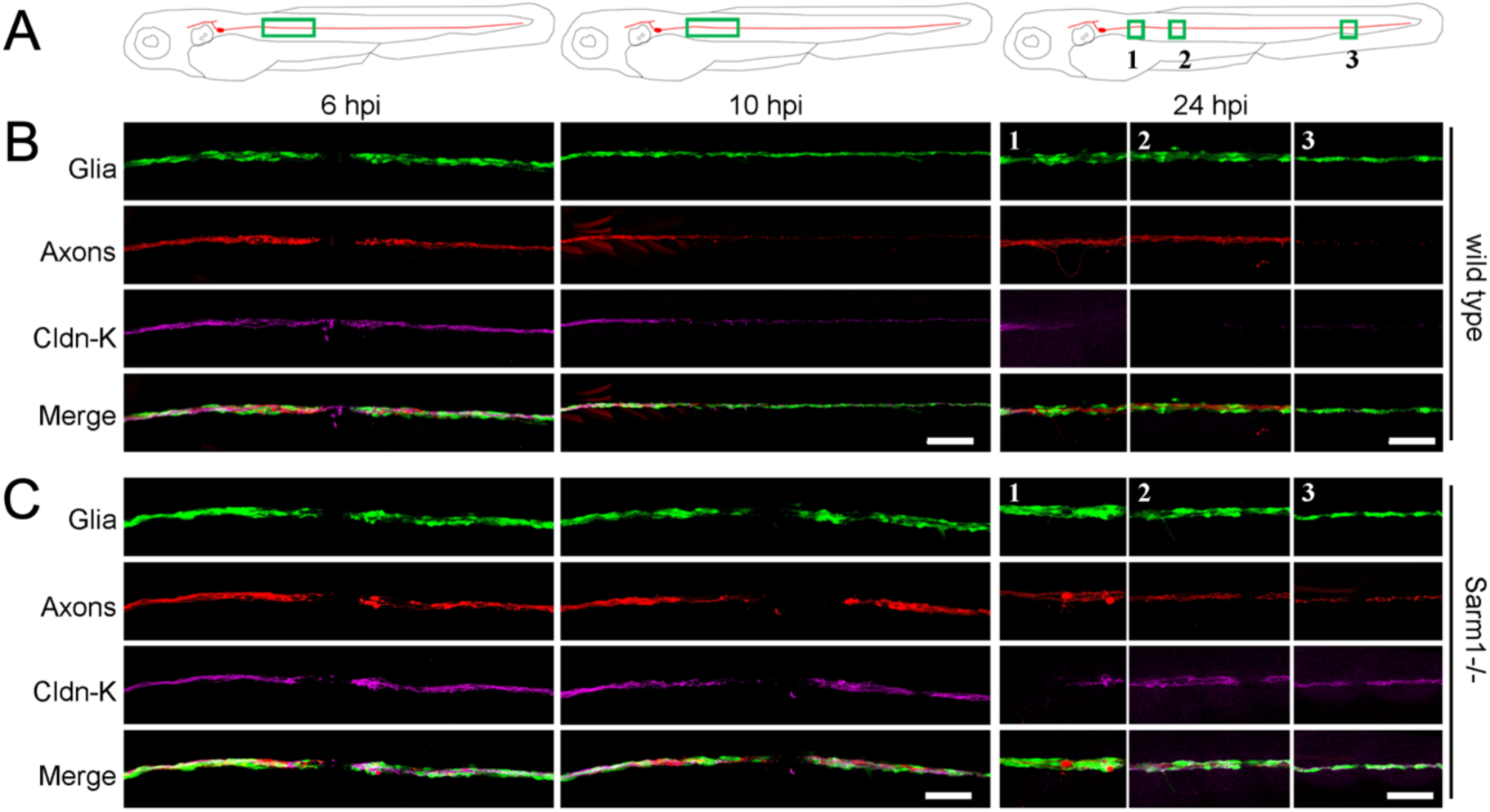
**A-C)** Schematic model of the confocal imaging locations on severed axons (A). Confocal images of wild type (B) and Sarm1-/- (C) specimens in 6hpi, 10hpi and 24hpi. hpi = hour post injury. The specimens were the Tg[gSAGFF202A;UAS:EGFP; SILL:mCherry] lines and were stained with Claudin-k (magenta) antibody. Scale bar 50μm.

Next, we assessed myelination using the 6D2 monoclonal antibody, which recognizes a carbohydrate epitope in the piscine P0-like myelin glycoproteins IP1 and IP2 (72, 73). As with Claudin-k, 6D2 labeling faded in Schwann cells distal to the injury in wild-type specimens (Figure 6A). Yet, 6D2 labeling remained unchanged in Sarm1 mutants (Figure 6B). We obtained congruent results when addressing myelination directly in living specimens by using a transgenic line expressing membrane-targeted EGFP under the control of the myelin binding protein (Mbp:EGFP-CAAX, green) (Figure 6C-D) (74). Together, these results indicate that Sarm1-deficient sensory axons are maintained independently of Schwann-cell support, and that the clearance the severed axons is not necessary for regenerating axon growth, pathfinding, myelination and re-innervation of sensory organs. In addition, they reveal that Schwann cells distal to the injury do not de-differentiate in Sarm1-mutant specimens.

**Figure 6.**
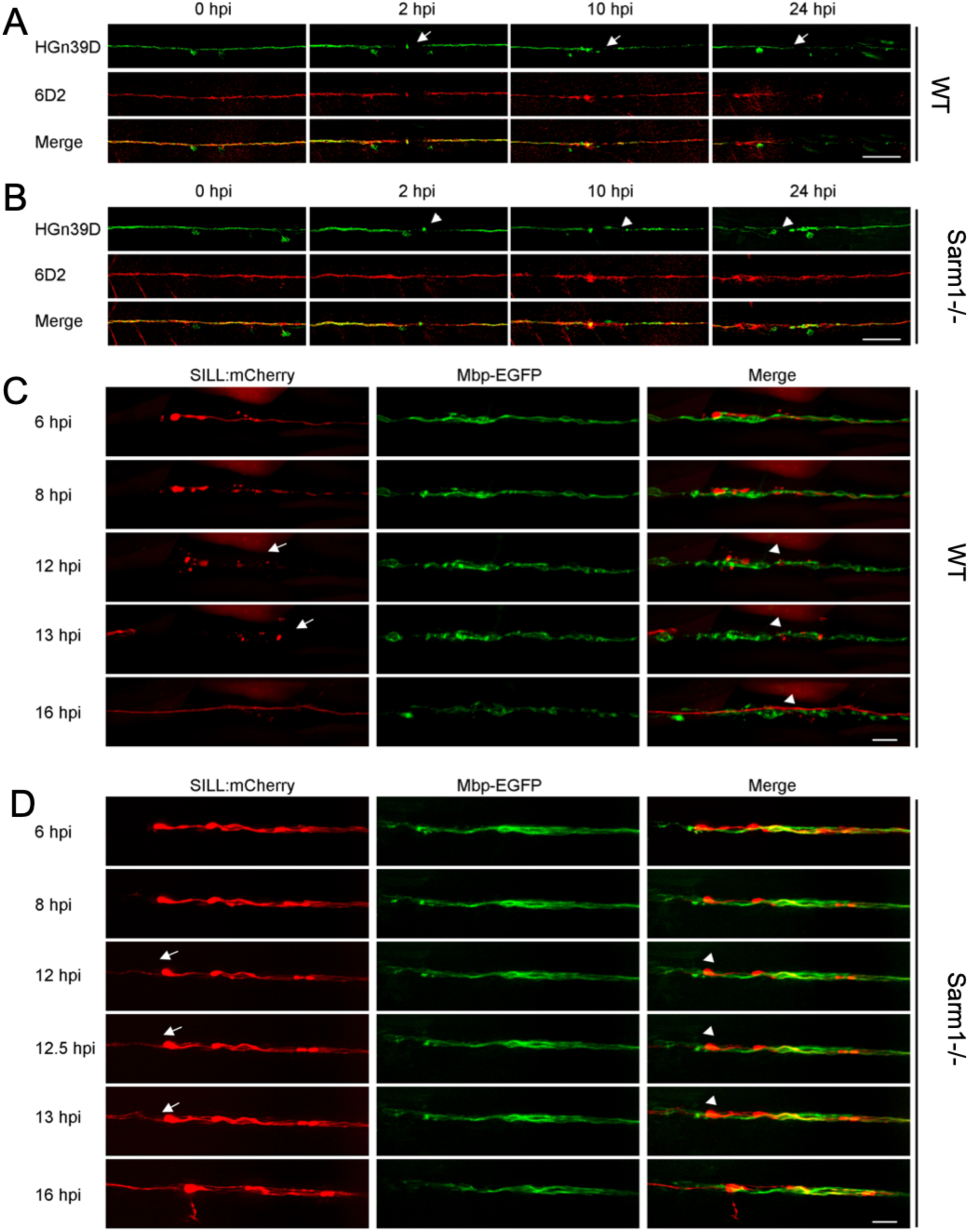
**A-B)** Confocal images of wild type (A) and Sarm1-/- (B) specimens expressing EGFP in sensory neurons of the lateral line (green) and stained with the monoclonal antibody 6D2. Stainings were performed at indicated time points after axons severing (hpi). The arrows point to the cutting sites. Scale bar is 50μm. **C)** Live imaging of the *Tg[Mbp-EGFP; SILL:mCherry]* after severing. The arrows indicated the fragmented axons and the arrowheads the fragmented myelin. Scale bar 20μm. **D)** Live imaging of the Sarm1-/- in *Tg[Mbp-EGFP; SILL:mCherry]* after severing. Arrows indicate regrowing axons, and arrowheads indicate the juxtaposition between the regrowing axons. Scale bar 20μm.

### Loss of Sarm1 counteracts Schwann-cell vulnerability to chemical toxins

Many chemotherapeutic agents invariably cause peripheral neurotoxicity, leading to permanent neuronal dysfunction (75–79). Under conditions of denervation, Schwann cells become strongly susceptible to chemotoxicity (80). We wondered if the protracted maintenance of severed axons in Sarm1 mutants may suppress glial vulnerability. To test this idea, we treated wild-type and Sarm1-mutant zebrafish with several chemical compounds that are under clinical trials or used as first-line treatment for common cancers in humans. First, we used 10-Hydroxycamptothecin (10-HCT), which is extremely toxic to denervated glia (82), and counted Schwann cells using the fluorescence transgenic marker *Tg[gSAGFF202A; UAS:EGFP]*, which is ideal for this experiment because the intensity of green fluorescence does not vary in denervated Schwann cells and, therefore, is independent of the maturity of these glia (41). We confirmed that 10-HCT does not affects in Schwann cells associated to viable axon (Figure 7A-B). However, it significantly reduced the number of Schwann cells in 10-HTC-treated wild-type animals after axon severing. By contrast, the number of distal Schwann cells was only marginally affected in Sarm1-mutant specimens (Figure 7B). Platinum-based, taxanes and some alkaloids are effective chemotherapeutic agents used as standards-of-care for various human malignancies, despite their severe neuropathic effects that include glial destruction (81). To address their effect on Schwann cell, we treated wild-type or Sarm1 mutant zebrafish with cisplatin, oxaliplatin, paclitaxel, docetaxel and vincristine. We found that upon nerve transection, all these drugs invariably killed injury-distal Schwann cells in wild-type specimens but not in Sarm1 mutants (Figure 7C). Importantly, none of these drugs affected Schwann cells associated with intact axons, suggesting that axons protect Schwann cells from chemical stress. To confirm this prediction, we forced the degradation of severed axons in Sarm1-deficient zebrafish treated with 10-HCT or vincristine. To this end, we repeated the used of capsaicin to activate rat TRPV1 expressed in a sub-set of lateralis neurons in homozygous mutant fish, in which Sarm1 is absent from every cell, including Schwann cells. This experiment revealed that Schwann cells lacking Sarm1 again become vulnerable to chemotoxicity once severed axons were synthetically eliminated (Figure 7D), confirming that Sarm1-mediated glioprotection is non-autonomous and depends upon the presence of non-degradable axons.

**Figure 7.**
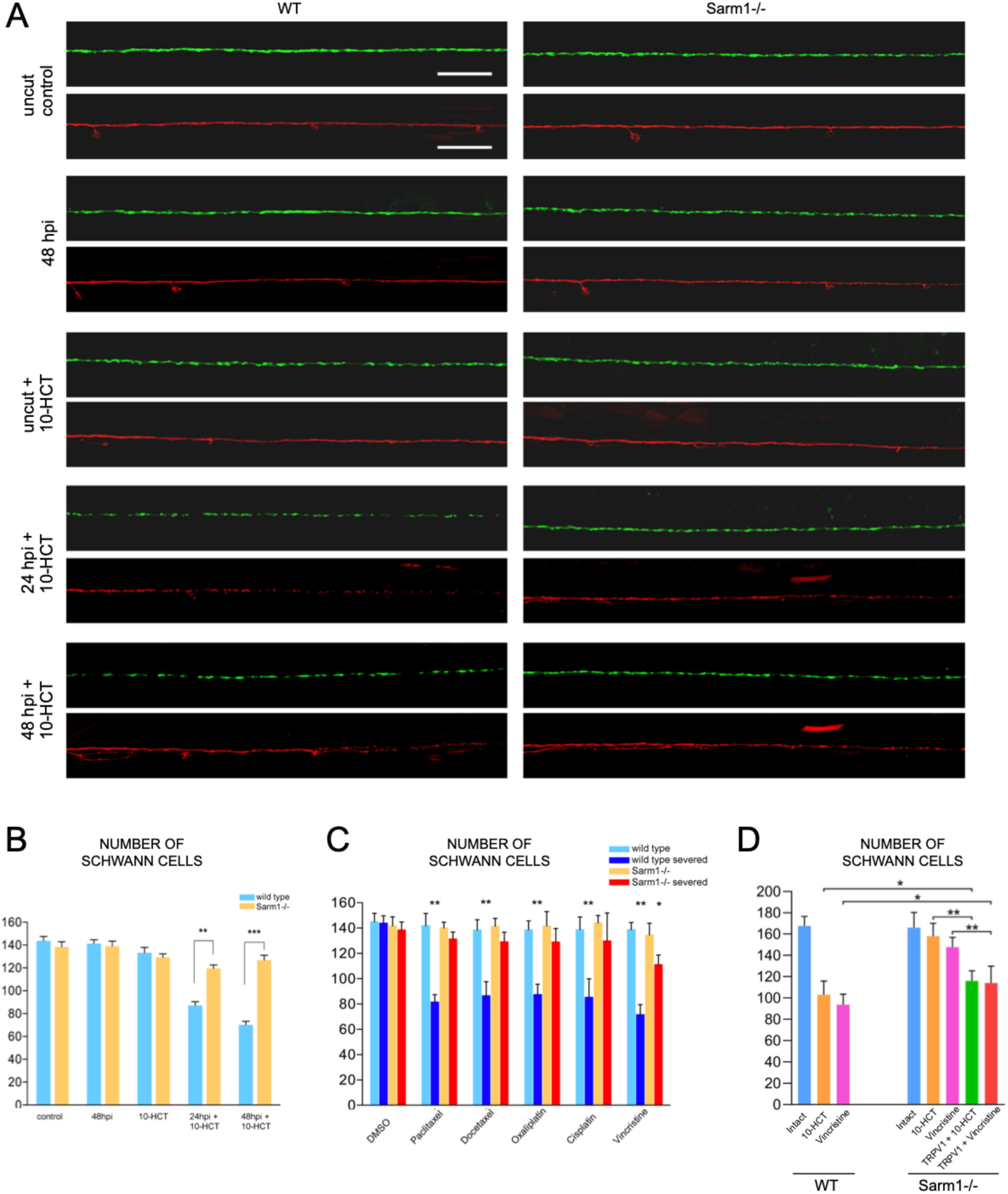
**A)** Confocal images showing Schwann cells (green) and lateralis sensory axons (red) in a control specimen (in which axons were not transected), in a specimen 48 hours after axon transection, and in specimens treated with 10-HCT (10-Hydroxycamptothecin). Left column is wild type and right column shows Sarm1-/-. In all cases, the concentration of 10-HCT in water was 40μm. Scale bar 100μm. **B)** Quantification of the Schwann cells from A). Data are shown as mean ± SEM. ** means p < 0.01, one-way ANOVA, n = 8 (each group). **C)** Quantification of Schwann cells of WT, WT severed, Sarm1-/- and Sarm1-/- severed with the treatment of the indicated chemical compounds for 48 hours. Concentrations: Paclitaxel 40μm, Docetaxel 0.1μm, Oxaliplatin 500μm, Cisplatin 50μm, Vincristine 50μm. Data are shown as mean ± SEM. * means p < 0.05; ** means p < 0.01, one-way ANOVA, n = 8 (each group). **D)** Quantification of Schwann cells after axon severing, in specimens treated with 10-HCT or Vincristine. Left bar group is wild type. Right bar group is Sarm1-/-, and Sarm1-/- with synthetically-eliminated axon segments.

## DISCUSSION

The sensory neurons that innervate skin, sensory receptors, joints and muscle communicate peripheral information to the brain, enabling animals to perform the essential activities of daily life. Nerve loss has significant negative impact on the quality of life of the affected individuals. Using zebrafish and a battery of tests that include subcellular structural characterization of sensory neurons and associated Schwann cells, neuronal-function, and behavioral assays, we offer a comprehensive and integrated analysis of neuronal and glial response to injury, as well as on the consequences of blocking axon degeneration systemically. It has been well established that the absence of Sarm1 in *Drosophila* and the mouse prevents the degradation of damaged axons (15, 83). Although we were not predicting any differences for Sarm1 in zebrafish, assaying functional conservation in our experimental neuronal pathway is not dispensable because some neurotraumatic conditions had been shown to lead to neuronal loss in the absence of Sarm1. Loss of Sarm1 improves functional recovery after traumatic brain injury in mice, and inhibits vincristine-mediated neurotoxicity (84, 85). Sarm1-deficient mice are viable (19, 86).We found that chronic and systemic loss of Sarm1 is compatible with zebrafish viability and sensorineural function, and demonstrate that the long-term maintenance of non-degradable axon fragments has no detrimental effect on the repair of a sensory circuit. Furthermore, regenerating axons fasciculate and myelinate normally, indicating that they do not compete with non-degradable axons segments for exiting glia. Therefore, we conclude that the axonal destructive and nerve reconstructive processes occur in parallel. In addition, these data suggest that a competitive balance between axon degradation and regeneration does not appear to have shaped the evolution of Sarm1 and, by extension, Wallerian degeneration.

Focusing on Schwann cells, we show that glial cells adjacent to nerve injury in Sarm1 mutants behave dramatically different than those of wild-type specimens. Specifically, Schwann cells in Sarm1 mutants do not migrate or extend projections to bridge the gap in the glial scaffold, indicating that these cells do not directly sense missing intercellular contacts or glial discontinuity. Instead, our findings suggest alternative scenarios. One is that the degradation of axons releases signals that induce Schwann cell to change behavior. In mice, for example, nerve damage promotes mesenchymal behavior of Schwann cells surrounding the wound via TGF-β signaling, which drives collective Schwann-cell migration across the wound (87). Although we did not observe Schwann-cell migration, signals derived from injured axons may promote wound-adjacent Schwann cells to extend projections to bridge the gap in a similar manner. Interestingly, we observed that filopodia-like structures emerged from Schwann cells at both sides of the injury in wild-type animals, suggesting that repair-inducing signals are likely diffusible, affecting glial cells independently of their association with axons. It remains to be determined if the source of such signals is the damaged axons, the denervated Schwann cells, or other cells in the wound microenvironment.

Upon nerve injury, activated macrophages rapidly accumulate around the wound and contribute to Wallerian degeneration and to axonal regeneration. Specifically, classical pro-inflammatory M1-type phagocytic macrophages remove axonal and myelin debris, whereas anti-inflammatory M2-type macrophages modulate Schwann-cell activity and promote axon regeneration. Immediately after nerve injury, Schwann cells release chemokines that attract or retain macrophages at the wound before axon fragmentation. Importantly, pharmacological activation of macrophage recruitment to the wound microenvironment enhances axon regeneration, whereas decrease of macrophage infiltration inhibits axon regeneration. Reactive oxygen species (ROS) are conserved early wound signals across species. Injury-mediated ROS production recruits inflammatory cells, including macrophages. Primary sources of ROS after nerve damage are the mitochondria and cellular NADPH oxidases. Studies in *ex vivo* cultured neurons indicate that Sarm1 acts downstream of mitochondrial ROS generation. Also, previous studies have characterized Wallerian axon degeneration and regeneration in Erbb2 mutant specimens, but did not address effects on the wound microenvironment and the behavior of macrophages (93–94). Accordingly, tested *in vivo* if the loss of Sarm1 as well as the absence of Schwann cells would affect macrophage recruitment, to find that the onset of recruitment, the retention time and number of macrophages at the wound did not differ between wild-type specimens and Sarm1 or Erbb2 mutants. Also, as in wild-type specimens, macrophages in Sarm1 and Erbb2 mutants engulfed debris locally, which were larger within macrophages in Ebbr2 mutants. Thus, the loss of Sarm1 does not affect focal damage resolution by macrophages, which are recruited and activated independently of the Schwann cells.

## CONCLUSIONS

Current mechanistic details about Sarm1 function derive from independent observations extrapolated from a wide variety of experimental systems. This has made it difficult to synthetize findings and reconcile some conflicting observations. Here, we have exploited a powerful *in vivo* genetic system to comprehensively study the consequence of systemic loss of Sarm1 from the sub-cellular to the organismal level in a single vertebrate system. By generating zebrafish carrying loss-of-function mutations in Sarm1, we confirmed and deepen previous findings from *Drosophila*, the mouse and cultured cells. Several novel insights derive from the work presented here. First, that loss of Sarm1 is well tolerated by the animal. Second, that neuronal-circuit repair is not contingent upon rapid clearance of damaged axons. Third, that Schwann cells are not necessary for maintenance of severed axons *in vivo*. Fourth, that after the axotomy Schwann cells distal to the cut site don’t dedifferentiate in Sarm1-mutants *in vivo*. Fifth, that the protracted maintenance of transected axons dramatically improves Schwann-cell tolerance to chemotoxicity after nerve injury after neurotrauma. This scenario is likely to occur in the clinical setting because nerve trauma is an inescapable consequence of surgical interventions, chronic metabolic dysfunction including diabetes, and pharmacological treatments such as antibiotics and anæsthesia that increase cellular stress (81). Therefore, these findings are of obvious pathophysiological significance. Crucially, these data lend strong support to the idea that direct interventions to systemically inhibit axon degradation are promising strategies to reduce chronic consequences of neurotrauma. Because TIR domain dimerization is necessary and sufficient to degrade NAD+, it renders Sarm1 amenable to inhibition by small molecules, as it has been demonstrated for the TIR domain of TLR2 (95). Our findings encourage the development of Sarm1 inhibitors for therapeutic applications (18, 29).

## MATERIALS & METHODS

### Zebrafish strains and husbandry

Zebrafish (*Danio rerio*) were maintained in a centralized facility in accordance to guidelines by the Ethical Committee of Animal Experimentation of the Helmholtz Zentrum München, the German Animal Welfare act Tierschutzgesetz §11, Abs. 1, Nr. 1, Haltungserlaubnis, to European Union animal welfare, and to protocols number Gz.:55.2-1-54-2532-202-2014 and Gz.:55.2-2532.Vet_02-17-187 from the “Regierung von Oberbayern”, Germany. The transgenic lines Tg[UAS:EGFP], Tg[HGn39D] and Tg[SILL:mCherry] (67), Tg[gSAGFF202A] (41), Tg[UAS:GCaMP7a] (88), Tg[mbpa:tgRFP-CAAX]^tum102Tg^ (also known as Tg[MBP:tgRFP]) (89) and Erbb2 mutants (41, 59) have been previously published. The Tg[Sarm1-/-] was generated by CRISPR/Cas9 mediated mutagenesis.

### Sarm1 mutagenesis

We used CRISPR/Cas9-mediated genome modification to generate loss-of-function mutations in exon 1 of Sarm1. Cas9 mRNA and sgRNAs were co-injected into one-cell stage embryos. Cas9 mRNA was generated *in vitro* from PmeI linearized CAS9 vector (pMLM3613) using the Ambion^TM^ mMESSAGE mMACHINE T7 Kit. The mRNA was purified with the RNA easy kit (Qiagen). To generate the sgRNAs, the target exon was sequenced and the sequence information was used to design oligonucleotides for the sgRNA guide vector (pDR274) using the on-line tool “ZiFiT Targeter software package” (http://zifit.partners.org/) (90). The sgRNA sequence for exon 1 of Sarm1 is 5’-GGGACTTGGAAGAGACCCGC-3’. The annealed oligonucleotide was cloned into the BsaI-digested pDR274 vector using T4 ligase (NEB M0202). Resulting clones were sequenced to verify correctness, and then linearized with DraI. The purified linearized DNA fragment was employed to generate the sgRNA using the T7 MEGAscript kit (Ambion^TM^). By out-crossing adult fish resulting from injection, we obtained germ-line transmission of two independent alleles: *sarm1^hzm13^* and *sarm1^hzm14^*. For genotyping mutant carriers, we used the primers: Forward: 5’-GATTTGCCGTTATCTCTCCA-3’ and Reverse: 5’-TCAAGCAGTTTGGCAGACTC-3’.

### DNA constructs

The DNA constructs SILL:mCherry, SILL:Gal4 (67), UAS:Synapsin1-GFP (53) and mbp:EGFP-CAAX have been previously described. The vectors pMLM3613 (42251) and pDR272 (42250) were purchased from Addgene. The plasmids UAS:TRPV1-tagRFP, the coding sequence of rat TRPV1 containing the E600K mutation fused to tagRFP was commercially synthesized by Genecat and the expression construct was generated using Tol2 kit. The constructs SILL:mito-mCherry, SILL:Sarm1-v2a-mCherry, SILL:Kaede, SILL:mitoRGECO and SILL:erGCaMP3 were generated using Tol2 kit. Mitochondria in lateralis neurons were marked by expressing the mitochondria targeting sequence from the Cytochrome-C oxidase subunit 8A fused to mCherry. The plasmids containing mito-RGECO and er-GCaMP3 (92) were a gift of David W. Raible (University of Washington).

### Antibodies and immunostaining

Whole-mount immunostaining was performed as following. First, samples of zebrafish embryos or larvae were fixed by immersion in ice-cold 4% paraformaldehyde diluted into phosphate-buffered saline buffer containing 0.2% Tween20 (PBST), and incubated overnight at 4°C. Then, the samples were washed at room temperature (RT) with PBST three times, 10 minutes per wash, and subsequently blocked in 10% bovine serum albumin (BSA), also at RT for 1 hour. Next, the samples were incubated in primary antibodies at 4°C overnight. Next, the samples were washed in PBST for 2 hours, changing to fresh buffer every 30 minutes. Finally, they were incubated with secondary antibodies at 4°C overnight. Primary antibodies and concentrations: mouse anti-Acetylated tubulin, 1:1000 (Sigma T7251); rat anti-Claudin-k, 1:500 (gift from T. Becker, University of Edinburgh, U.K.) (71); mouse 6D2, 1:5 (gift from Dr. G. Jeserich, University of Osnabrück, Germany) (73). Secondary antibodies used were at the following concentrations: donkey anti-Mouse Alexa Fluor® 555, 1:200, Abcam ab150106; donkey anti-Rat IgG H&L (Alexa Fluor® 647) pre-adsorbed, 1:200, Abcam ab150155). Samples were washed in PBST for 30 minutes and mounted in Vectashield one day before microscopic examination. Imaging of fixed samples was done with a laser-scanning confocal microscope (LSM 510, Carl Zeiss).

### Intravital microscopy

For videomicroscopy, larvae were anesthetized with MS-222 (0.013% M/V) in Danieau’s and mounted in 0.8% low melting-point agarose on a 35mm glass-bottom Petri dish. Samples were gently pressed against the glass using a hair-loop glued to the tip of a glass pipette, as previously described. The agarose dome was immersed in Danieau’s with the anesthetic. Images of the axons and Schwann cells were acquired using a spinning-disc microscope with a 40× air objective at 28.5°C (58). Z-stacks were set to 0.8-1.2 µm intervals. Time intervals were 10 minute or 15 minute per stack. The resulting raw data were processed, assembled and analyzed with ImageJ.

### Western blot assay

Wild-type and mutant larvae were anesthetized and killed 5 days-post-fertilization. Samples were homogenized in ice-cold RIPA buffer (with protease inhibitor cocktail from Roche (Cat.04693159001). After homogenization, the samples were incubated on ice for 30 minutes for further lysis. The resulting lysate was centrifuged at 1200 rpm at 4°C, and the supernatant was taken for the BCA assay. The supernatant was diluted in loading buffer and boiled at 99°C for 5 minutes. Next, the samples were performed in the SDS-PAGE and transferred onto a PVDF membrane. After blocking the membrane in 5% skimmed milk (diluted in PBST) for 1 hour, the membrane was incubated with the primary antibody (rabbit anti-Sarm1, 1:500, ANASPEC 55381; Mouse anti-β-Tubulin, 1:2000, Sigma T5168) at 4°C overnight. The next day, the membrane was washed and incubated with HRP-labeled secondary antibody (Peroxidase-Affini Pure Goat Anti-Mouse IgG (H+L), 1:10000, Jackson Immuno Research 115035003; Peroxidase-Affini Pure Goat Anti-Rabbit IgG (H+L), 1:10000, Jackson ImmunoResearch 115035144) for 1 hour. Images were acquired by developing the membrane with ECL (Pierce™ ECL Western Blotting Substrate, Thermo Fisher, 32109).

### Laser microsurgery

To mark lateralis sensory neurons individually, DNA of the SILL:mCherry construct was injected into eggs of Tg[HGn39D], Tg[HGn39D; Sarm1-/-], Tg[SILL:mCherry; gSAGFF202A; UAS:EGFP] or Tg[SILL:mCherry; gSAGFF202A; UAS:EGFP; Sarm1-/-] zebrafish. Resulting larvae were selected according to red-fluorescence in lateralis neurons. Selected samples were mounted into agarose as described above, and their peripheral axons were targeted an ultraviolet laser (350nm) using the iLasPulse system (Roper scientific AS, Evry, France), as described previously. The laser beam was delivered using a 63x water-immersion objective (50, 53, 58). The laser pulses were calibrated and applied to the target area until a clear gap in the axons was visible. The samples were observed again 1 hour post-injury (hpi) to confirm that the axons were completely transected.

### Quantification of mitochondrial density and motility

To analyze mitochondria in sensory axons, we generated kymographs of mito-mCherry fluorescent spots using the Multi-Kymograph tool of the Fiji software (http://fiji.sc). The movement of mitochondria was determined by the slope of the lines drawn over time, and the direction of movement was determined by the moving mitochondria crossing the time line (vertical) in the kymographs. The data were analyzed with Python scripts and the Graphpad Prism software.

### Calcium imaging

For calcium imaging in lateralis neurons, Tg[SILL:Gal4; UAS:GCaMP7a] double-transgenic larvae were anesthetized and mounted in 0.8% low melting-point agarose on a 35mm glass-bottom Petri dish. Imaging was acquired through a 63x water-immersion objective with an exposure time of 400 milliseconds. Laser-mediated axon transection was done after the 4^th^ imaging of the time series. Next, live videomicroscopy was done for 2 minutes at a frame rate of 400 milliseconds at 28.5°C. The raw data were analyzed with ImageJ. To quantify the calcium signal, the images were processed to ImageJ. The region of interest (ROI) was selected and measured the value with time point. GCaMP or RGECO intensity changes were calculated as follows: ΔF/F0 = (F–F0)/F0, where F0 is the value of the fluorescent signal before axons were transected, and F is the value of the fluorescent signal with time point after axon severing (92).

### Chemogenetics

For chemogenetic experiments, we co-injected the SILL:Gal4 with UAS:mCherry or UAS:ratTRPV1-tagRFP into the Tg[UAS:GCaMP7a] or Tg[HGn39D; Sarm1-/-]. The positive larva with SILL:Gal4; UAS:ratTRPV1-tagRFP; UAS:GCaMP7a expression was used to activate TRPV1 channels in zebrafish by incubation in 5μM capsaicin (Sigma, M2028) and subsequent live imaging for 1hour of the mounted and anesthetized embryo. Images were acquired through a 63x water-immersion objective with an exposure time of 400 milliseconds. For experiments with the Sarm1 mutant larvae, the HGn39D with SILL:mCherry or ratTRPV1-tagRFP positive animals were laser axotomized. The larvae were treated with 10 μM capsaicin or ethanol (1:1000, v/v) 2h after transection. 1.5h after capsaicin treatment, images were taken by spinning disc microscopy.

### Behavioral assays

For the touch-mediated escape response, 2dpf embryos were gently dechorionated and kept in Danieau’s solution at 28°C for at least 1 hour. Embryos were place into a flat uncovered Petri dish containing Danieau’s and were recorded with a high-speed camera (NX4 series, Imaging solution, GmbH). Video recording was launched and a randomly chosen embryo was touched with a blunt glass needle until it evoked a reaction. Recordings were done under white-light illumination over 150 seconds at a rate of 200 frames per second (fps). The swimming trajectories were obtained with 3D Particle Tracker plugin, ImageJ software. The further quantification and statistics were using Python.

### Statistical analysis

The Student’s t test (two tailed), one-way ANOVA test and Wilcoxon rank sum test were applied using Python scripts. Error bars in all figures are standard errors of the mean (SEM).

## Acknowledgements

Acknowledgements

This research was supported by a Grant from the Human Frontiers Science Programme and by the Helmholtz Gemeinschaft to H.L-S.

## SUPPLEMENTAL MATERIALS

**Supplemental Figure 1.**
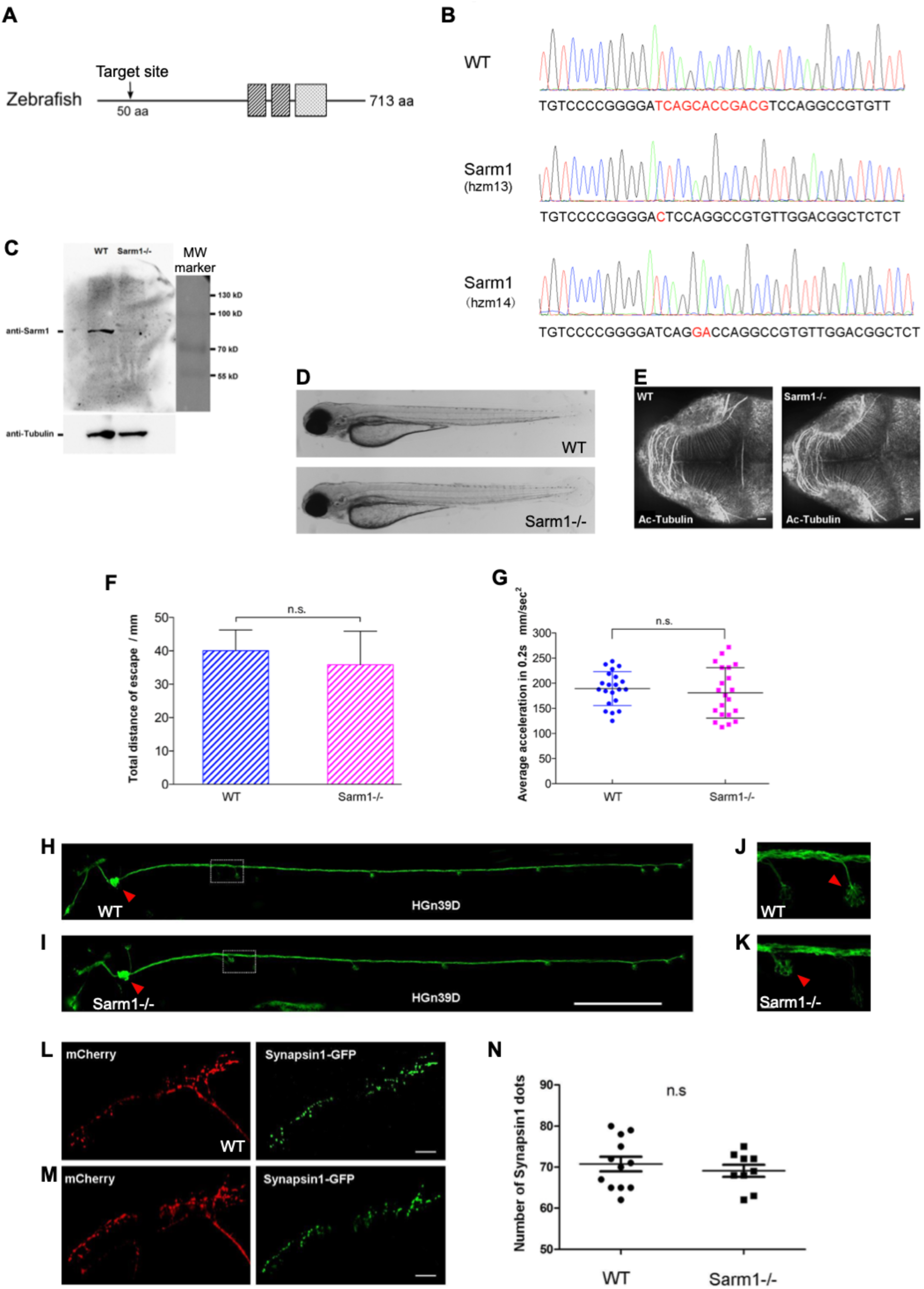
**A)** Structure of the zebrafish Sarm1 protein highlighting the two SAM domains (dark grey), the TIR domain (light grey). The downward arrow indicated the approximate region targeted for mutagenesis, approximately 50 amino acids from the start codon. **B)** Sequence of the wild-type Sarm1 indicating in red the mutagenized area and, below, the two mutant alleles obtained in this study. The *hzm13* allele introduces an 11-base deletion and T/C mutation, resulting in a frame shift and premature stop codon. The *hzm14* allele is a 7-base deletion and AG/GA mutation that also generates a frame shift and premature stop codon. **C)** Western blot of protein extracts from wild type and Sarm1*^hzm13^* fish embryos using a commercial ant-Sarm1 antibody, revealing absence of the protein in the mutants. An antibody to alpha-Tubulin was used as loading control. **D)** Low-magnification image of a wild-type 5dpf zebrafish (top) and a homozygous Sarm1*^hzm13^* (bottom), showing no overall anatomical differences. **E)** Confocal image of a wild-type 5dpf zebrafish (left) and a homozygous Sarm1*^hzm13^* (right) stained with an antibody to acetylated Tubulin to mark neurons in the central nervous system, showing no evident defects in the mutants. In this and all figures, rostral is left and caudal is right. Scale bar 20µm. **F-G)** Quantification of sensorimotor function in zebrafish. (H) shows the total distance traveled by larvae after touch-trigger escape response in wild-type (dashed blue bar) and homozygous Sarm1*^hzm13^* (dashed magenta bar). (I) dot plot of the average acceleration of wild-type (blue) and homozygous Sarm1*^hzm13^* (magenta) after tactile touch-induced escape response. Error bar = SEM; n.s. means no significant difference, Student’s t test. wild type n=21, Sarm1-/- n=21. **H-I)** Confocal image of a 5dpf wild-type (H) and Sarm1-/- (I) larvae carrying the *Tg[HGn39D]* transgene to mark lateralis afferent neurons with GFP. The posterior lateral-line ganglion is indicated with a red arrowhead. The dotted box indicates an innervated neuromast (expanded in C). Scale bar 400µm. **J-K)** Confocal image of the peripheral arborization of lateralis neurons in 5dpf wild-type (H) and Sarm1-/- (I). Red arrows indicate the position of a neuromast from the dotted boxes in (H-I). **L-M)** Confocal image of the central arborization of lateralis neurons in 5dpf wild-type (L) and Sarm1-/- (M), mCherry (red) and Synapsin1-GFP (green) to reveal normal arborization and pre-synaptic puncta in both cases. **N)** Quantification of the number of synapsin1 puncta from (E-F), Error bar = SEM; n.s. = not significant, Student’s t test. Wild type n=15, Sarm1-/- n=15.

**Supplemental Figure 2.**
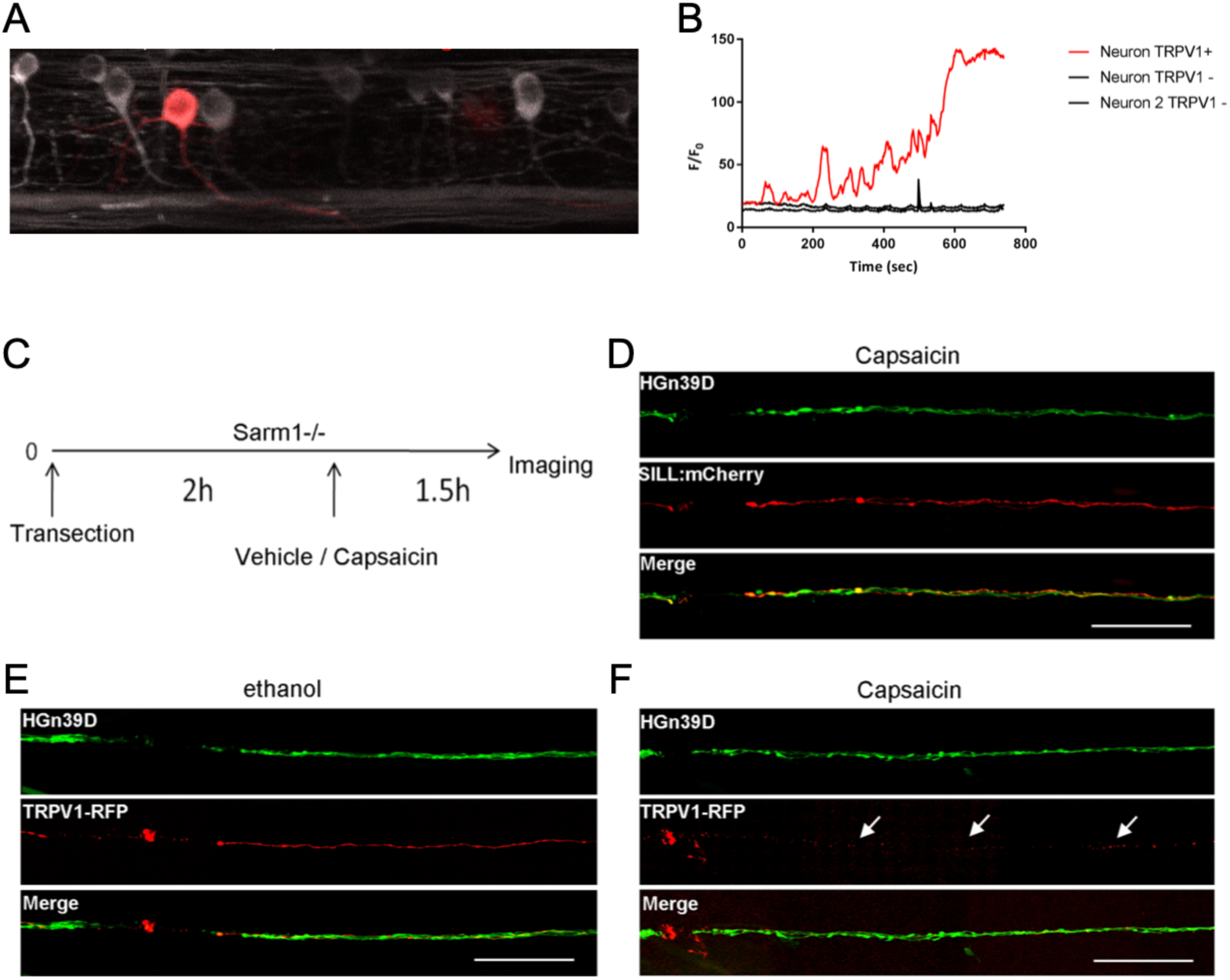
**A)** Confocal image of UAS-TRPV1-tagRFP1 (red, injected) and UAS-GCaMP (green, transgenic) labeled neurons with Cntn1b:KalTA4. **B)** Quantification of the calcium signal after capsaicin incubation. Red plot indicated the TRPV1 positive neuron. The black plots indicated two TRPV1 negative neurons. **C)** Schematic representation of the experimental strategy to synthetically elevate calcium in Sarm1-deficient transected axons. Lateralis sensory neurons were made to express a transgene coding for the rat transient receptor potential cation channel subfamily V member 1 (TRPV1) fused to RFP, or simply mCherry. Two hours after axon transection, zebrafish larvae were bathed in ethanol solution (control), or ethanol containing capsaicin, a natural activator of TRPV1. 90 minutes after treatments, larvae were imaged by confocal microscopy to assess the extent of distal segment degradation. **D)** Sarm1-/- fish expressing GFP in all lateralis neurons (*Tg[HGn39D]*) and mCherry in a mosaic manner in some neurons. Scale bars 100μm. **E)** Sarm1-mutant fish expressing GFP in all lateralis neurons and TRPV1-RFP in a mosaic manner. Scale bars 100μm. **F)** Sarm1-mutant fish expressing GFP in all lateralis neurons and TRPV1-RFP in a mosaic manner. Capsaicin treatment induced transected axon degradation (former location of the axon signaled by three white arrows). Scale bars 100μm.

**Supplemental Figure 3.**
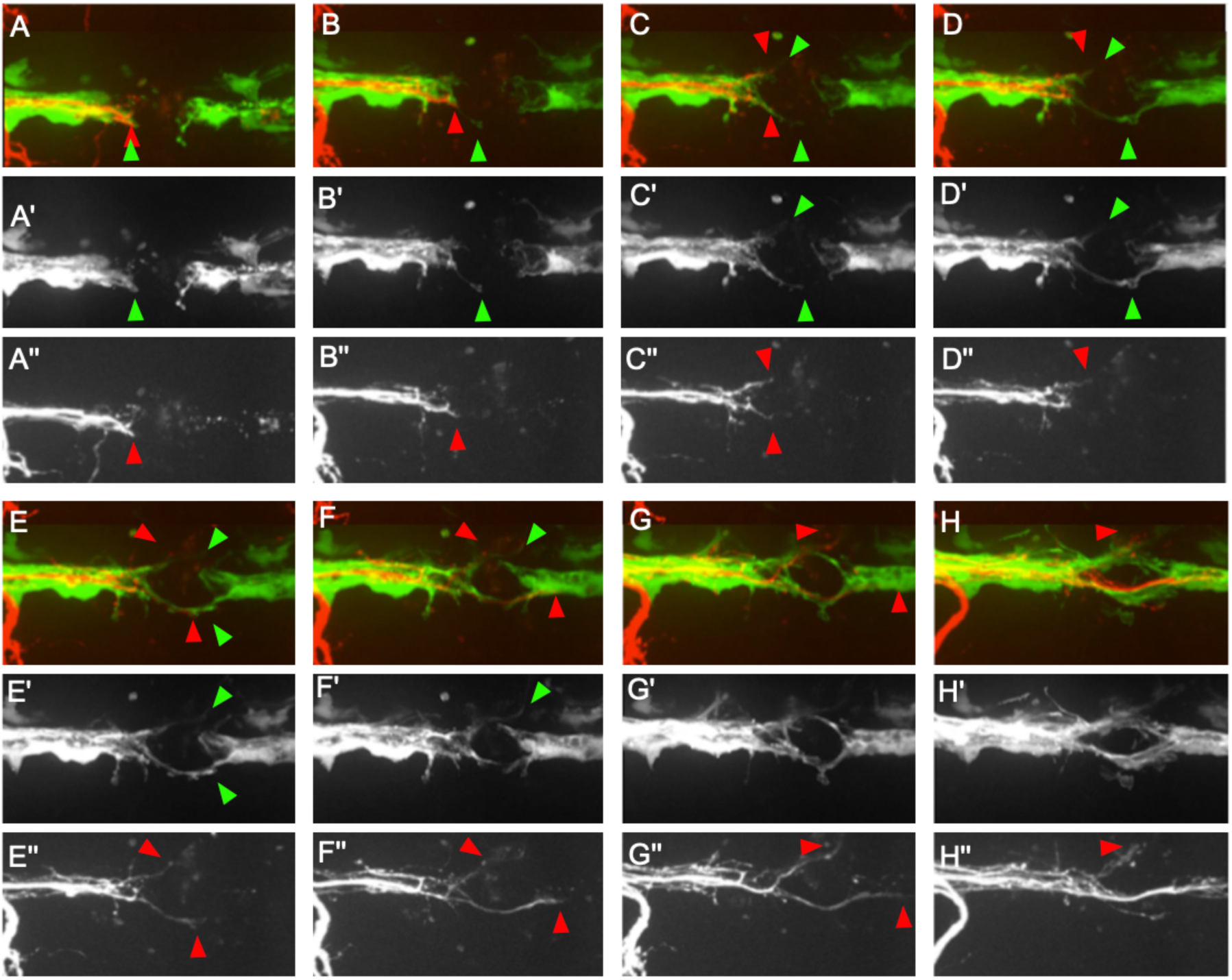
Series of still images taken from Supplemental Movie 1 (also shown in Figure 6, panel E). They show eight time points during the repair of the gap in the glial scaffold (A’-H’) and axonal regeneration after transection (A’’-H’’) in a wild-type animal. Rostral is left and caudal is right. In all panels, the red arrowheads signal the location of the pioneering growth cone of the regenerating axons. The green arrowheads mark the filopodia-like extensions from Schwann cells adjacent to the glial gap. At the start of the series, the axonal terminal stump and the Schwann cells proximal to the gap co-localize (juxtaposition of the green and red arrowhead). In C’, a Schwann cell extends a filopodium across the gap, whereas the axons (C’’) do not grow along this extension of across the gap. In D’, several extensions from Schwann cells are clearly visible at the top and bottom aspects of the image. The shape of the lower extension from a Schwann cell did not change shape, suggesting that they are stabilized, perhaps through interactions with the substrate. Proximal axons (D’’) start to grow along these Schwann-cell protrusions. In E’, the extensions from the anterior and posterior Schwann cells have crossed the gap, physically interact and commence to reconstitute a continuous glial scaffold. The axonal projections (E’’), however, have suffered a retraction towards the proximal stump. F’-H’ show a continuation of Schwann cells behavior, increasing the contacts and closing the gap, which is obvious by the smaller area of the gap (distance between proximal and distal Schwann cells). F’’-H’’ show a more robust and persistent extension of the axonal growth cones, which grow nearly strictly along the Schwann cells extensions. Although the gap in the glial scaffold is much reduced, nerve fibers show discrete defasciculation (H’’).

**Supplemental Figure 4.**
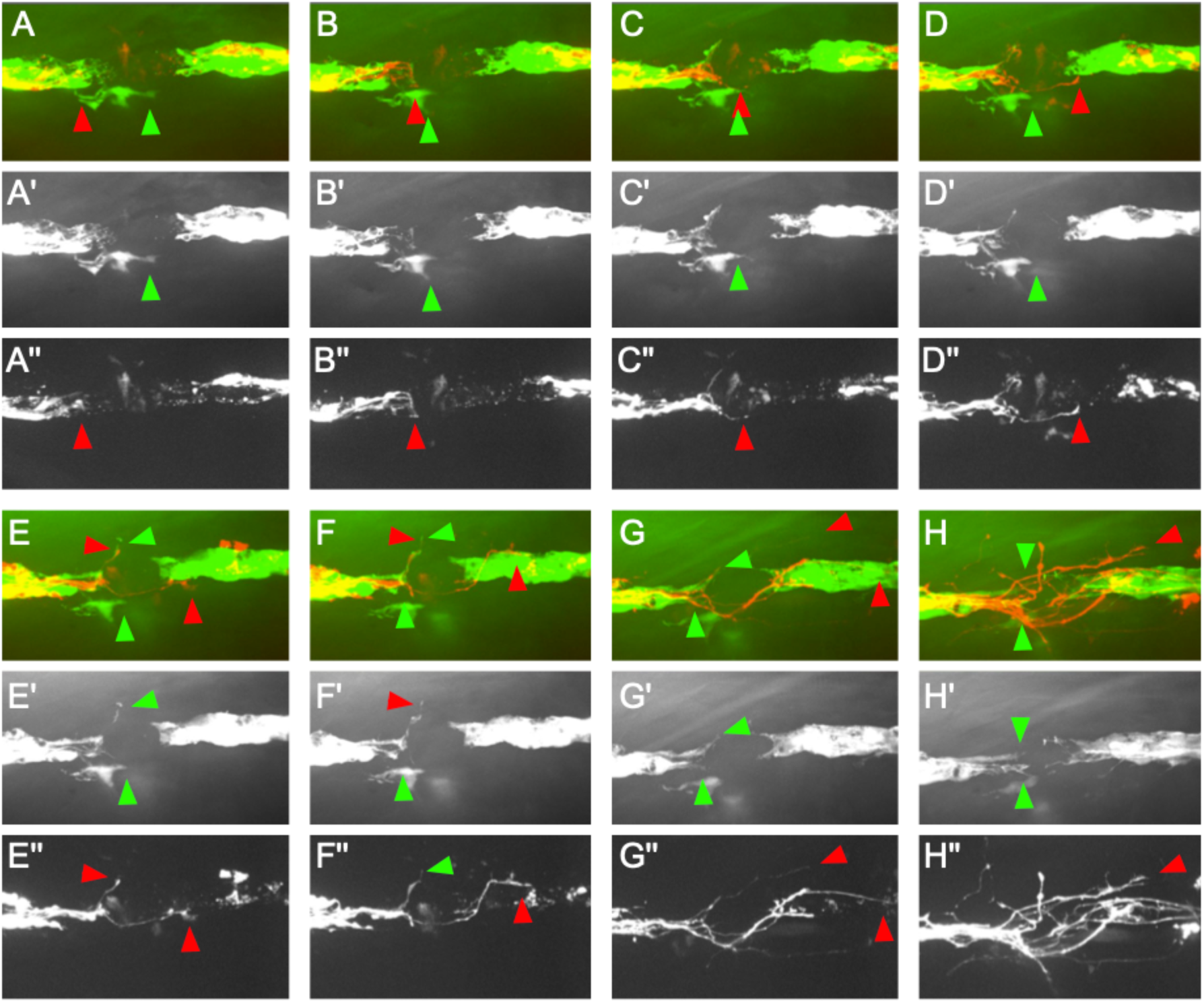
Series of still images taken from Supplemental Movie 2 (also shown in Figure 6, panel F). They show eight time points during the repair of the gap in the Schwann-cell scaffold (A’-H’) and axonal regeneration after transection (A’’-H’’), in a Sarm1-mutant specimen. In all panels, the red arrowheads signal the location of the pioneering growth cone of the regenerating axons, and the green arrowheads mark filopodia-like extensions from Schwann cells. Unlike the wild-type situation shown in Supplemental Figure 1, the Schwann cells adjacent to the gap form small filopodia-like extensions, but which never cross the gap. The proximal axon stumps eventually form growth cones that cross the gap at various locations and, upon finding distal Schwann cells, grow along the glial scaffold (E’’-H’’). Nerve fibers show extensive local defasciculation (H’’).

**Supplemental Movie 1.** Videomicroscopic imaging of a double transgenic *Tg[SILL:mCherry; Mfap4-memEGFP]* 4dpf wild-type specimen in which the mCherry(+) axons were severed by a laser pulse. The video represents for 13 hours of continuous imaging at a 5-minute temporal resolution. Axons (red) and macrophages (green) were visualized relative to the cut (proximal is on the left and distal in on the right).

**Supplemental Movie 2.** Videomicroscopic imaging of a double transgenic *Tg[SILL:mCherry; Mfap4-memEGFP]* 4dpf specimen that was mutant for Sarm1, in which the mCherry(+) axons were severed by a laser pulse. The video represents for 13 hours of continuous imaging at a 5-minute temporal resolution. Axons (red) and macrophages (green) were visualized relative to the cut (proximal is on the left and distal in on the right).

**Supplemental Movie 3.** Videomicroscopic imaging of a double transgenic *Tg[SILL:mCherry; Mfap4-memEGFP]* 4dpf specimen that was mutant for Erbb2, in which the mCherry(+) axons were severed by a laser pulse. The video represents for 13 hours of continuous imaging at a 5-minute temporal resolution. Axons (red) and macrophages (green) were visualized relative to the cut (proximal is on the left and distal in on the right).

**Supplemental Movie 4.** Videomicroscopic imaging of a 4dpf wild-type specimen carrying three transgenes Tg[SILL:mCherry; gSAGFF202A; UAS:EGFP], showing the dynamics of axons and Schwann cells after axonal transection. Sensory axons are shown in red and Schwann cells in green. The movie is a maximal projection of confocal stacks. It was recorded during 12 hours at 10-minute intervals. The top panel shows the merged images, the middle panel the Schwann cells, while the bottom panel the axons.

**Supplemental Movie 5.** Identical experiment as shown in Supplemental Movie 1, but conducted in a Sarm1-mutant specimen. It was recorded during 12 hours at 10-minute intervals. The top panel shows the merged images (SILL:mCherry; gSAGFF202A; UAS:EGFP), the middle panel the Schwann cells (gSAGFF202A; UAS:EGFP), and the bottom panel the axons (SILL:mCherry).

