## Supplementary material for "Loss of Sarm1 non-autonomously protects Schwann cells from chemotoxicity": Supp Figs

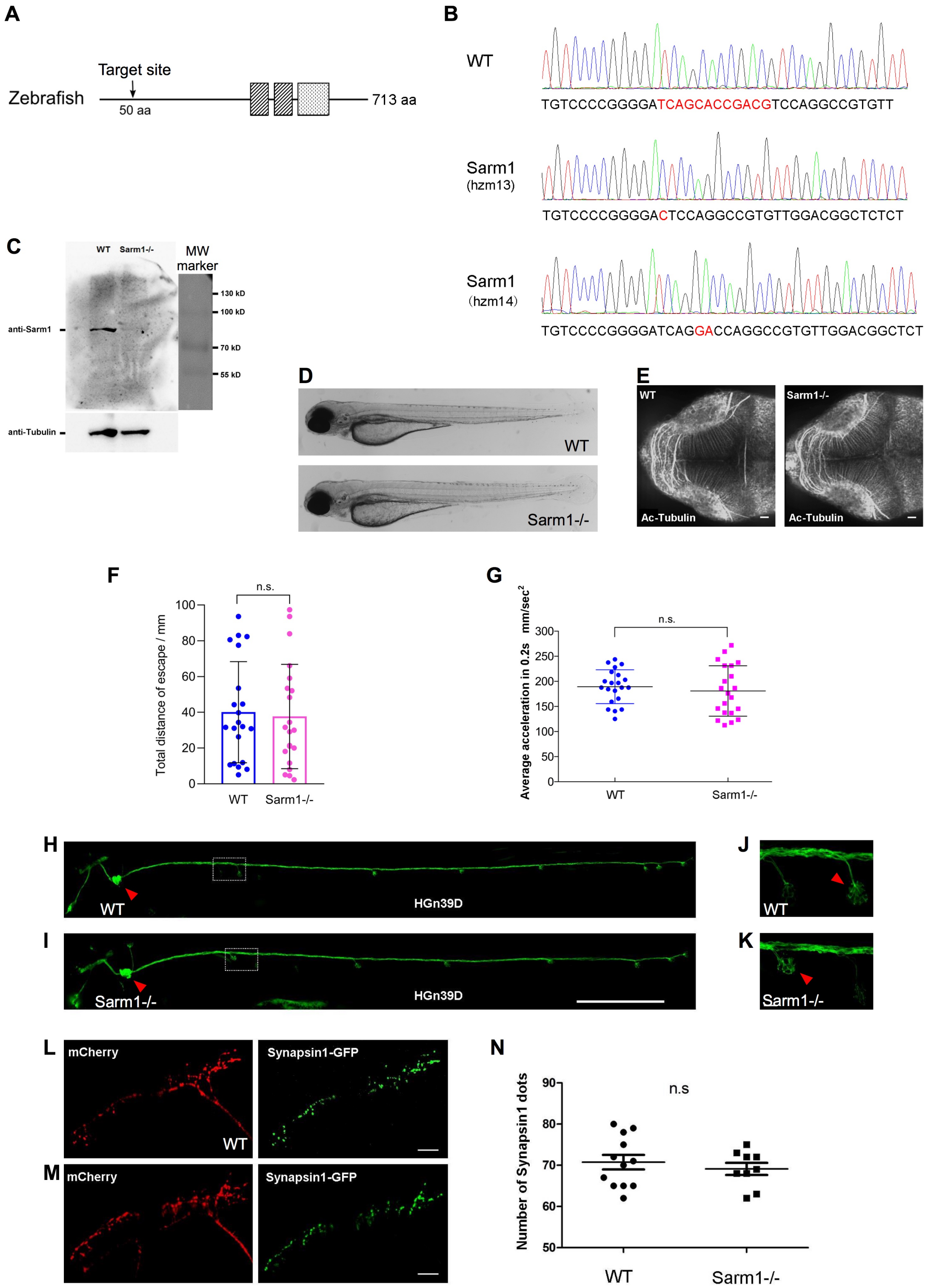


**Supplemental Figure 1. A)** Structure of zebrafish Sarm1 highlighting the two SAM domains (dark grey), the TIR domain (light grey). The downward arrow indicated the region targeted for mutagenesis, approximately 50 codons from the start codon. **B)** Sequence of the wild-type Sarm1 indicating in red the mutagenized area and, below, the two mutant alleles obtained in this study. The ***hzm13*** allele introduces an 11-base deletion and T/C mutation, resulting in a frame shift and premature stop codon. The ***hzm14*** allele is a 7-base deletion and AG/GA mutation that also generates a frame shift and premature stop codon. **C)** Western blot of protein extracts from wild type and Sarm1$^{hzm13}$ fish embryos using a commercial ant-Sarm1 antibody, revealing absence of the protein in the mutants. An antibody to alpha-Tubulin was used as loading control. **D)** Low-magnification image of a wild-type 5dpf zebrafish (top) and a homozygous Sarm1$^{hzm13}$ (bottom), showing no overall anatomical differences. **E)** Confocal image of a wild-type 5dpf zebrafish (left) and a homozygous Sarm1$^{hzm13}$ (right) stained with an antibody to acetylated Tubulin to mark neurons in the central nervous system, showing no evident defects in the mutants. In this and all figures, rostral is left and caudal is right. Scale bar 20µm. **F-G)** Quantification of sensorimotor function in zebrafish. (H) shows the total distance traveled by larvae after touch-trigger escape response in wild-type (dashed blue bar) and homozygous Sarm1$^{hzm13}$ (dashed magenta bar). (I) dot plot of the average acceleration of wild-type (blue) and homozygous Sarm1$^{hzm13}$ (magenta) after tactile touch-induced escape response. Error bar = SEM; n.s. means no significant difference, Student's t test. wild type n=21, Sarm1-/- n=21. **H-I)** Confocal image of a 5dpf wild-type (H) and Sarm1-/- (I) larvae carrying the ***Tg[HGn39D]*** transgene to mark lateralis afferent neurons with GFP. The posterior lateral-line ganglion is indicated with a red arrowhead. The dotted box indicates an innervated neuromast (expanded in C). Scale bar 400µm. **J-K)** Confocal image of the peripheral arborization of lateralis neurons in 5dpf wild-type (H) and Sarm1-/- (I). Red arrows indicate the position of a neuromast from the dotted boxes in (H-I). **L-M)** Confocal image of the central arborization of lateralis neurons in 5dpf wild-type (L) and Sarm1-/- (M), mCherry (red) and Synapsin1-GFP (green) to reveal normal arborization and pre-synaptic puncta in both cases. **N)** Quantification of the number of synapsin1 puncta from (E-F), Error bar = SEM; n.s. = not significant, Student's t test. Wild type n=15, Sarm1-/- n=15.

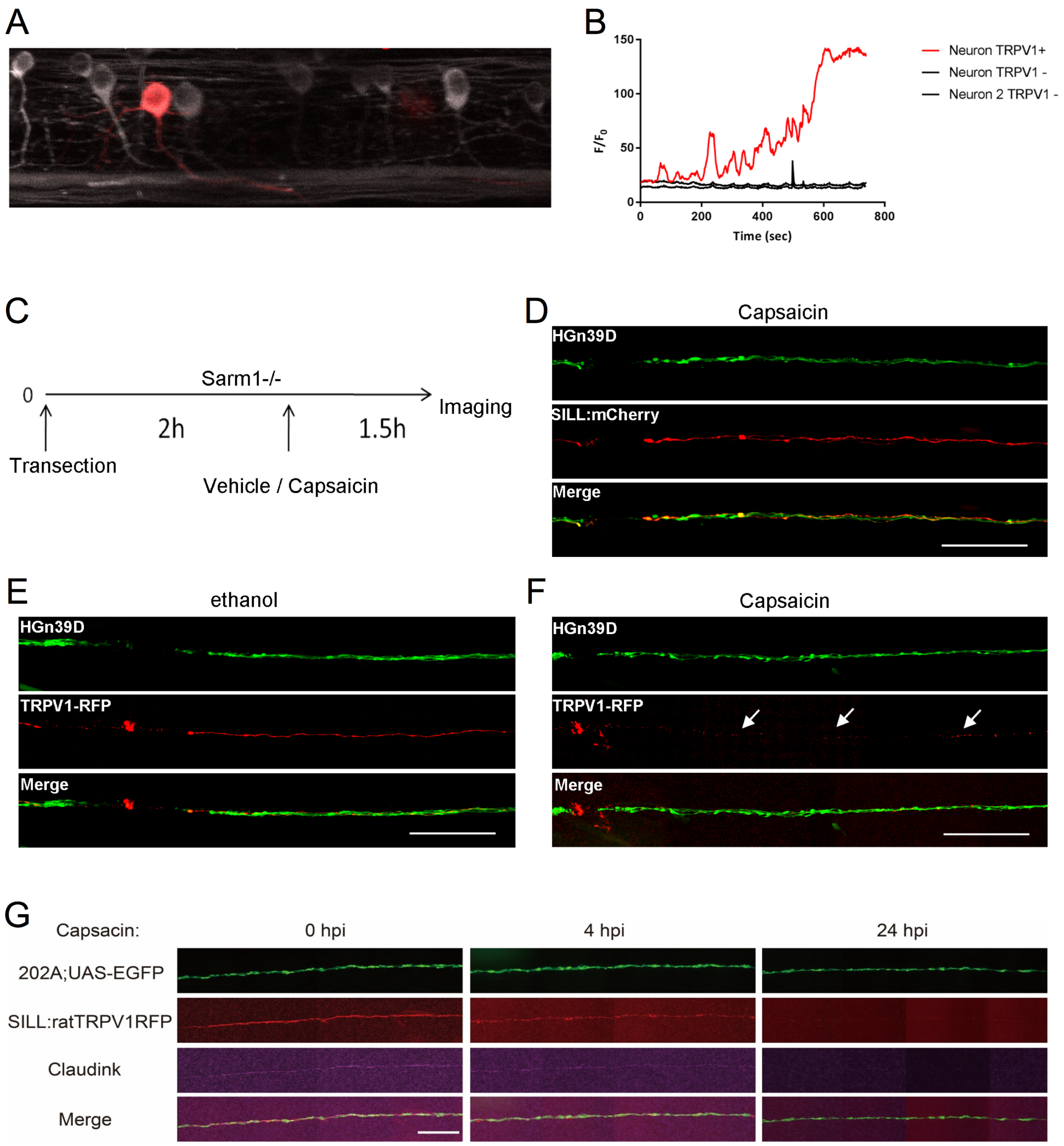


**Supplemental Figure 2. A)** Confocal image of UAS-TRPV1-tagRFP1 (red, injected) and UAS-GCaMP (green, transgenic) labeled neurons with Cntn1b:KalTA4. **B)** Quantification of the calcium signal after capsaicin incubation. Red plot indicated the TRPV1 positive neuron. The black plots indicated two TRPV1 negative neurons. **C)** Schematic representation of the experimental strategy to synthetically elevate calcium in Sarm1-deficient transected axons. Lateralis sensory neurons were made to express a transgene coding for the rat transient receptor potential cation channel subfamily V member 1 (TRPV1) fused to RFP, or simply mCherry. Two hours after axon transection, zebrafish larvae were bathed in ethanol solution (control), or ethanol containing capsaicin, a natural activator of TRPV1. 90 minutes after treatments, larvae were imaged by confocal microscopy to assess the extent of distal segment degradation. **D)** Sarm1-/- fish expressing GFP in all lateralis neurons (***Tg[HGn39D]***) and mCherry in a mosaic manner in some neurons. Scale bars 100µm. **E)** Sarm1-mutant fish expressing GFP in all lateralis neurons and TRPV1-RFP in a mosaic manner. Scale bars 100µm. **F)** Sarm1-mutant fish expressing GFP in all lateralis neurons and TRPV1-RFP in a mosaic manner. Capsaicin treatment induced transected axon degradation (former location of the axon signaled by three white arrows). Scale bars 100µm. PS: Tg[Sarm1-/-; 202A; UAS-EGFP; SILL:mCherry], 4dpf, incubated with capsaicin for 2 hours, then withdraw capsaicin. **G)** These images show a Sarm1-mutant specimen. Schwann cells (green) were stained with anti-Claudin-k antibody (magenta) and neurons expressing ratTRPV1-RFP (red), with indicated time points after axon severing and capsaicin treatment (hpi : hours post induction), showing that upon synthetic axonal degradation by TRPV1 activation, Schwann cells cease to express a terminal differentiation marker.

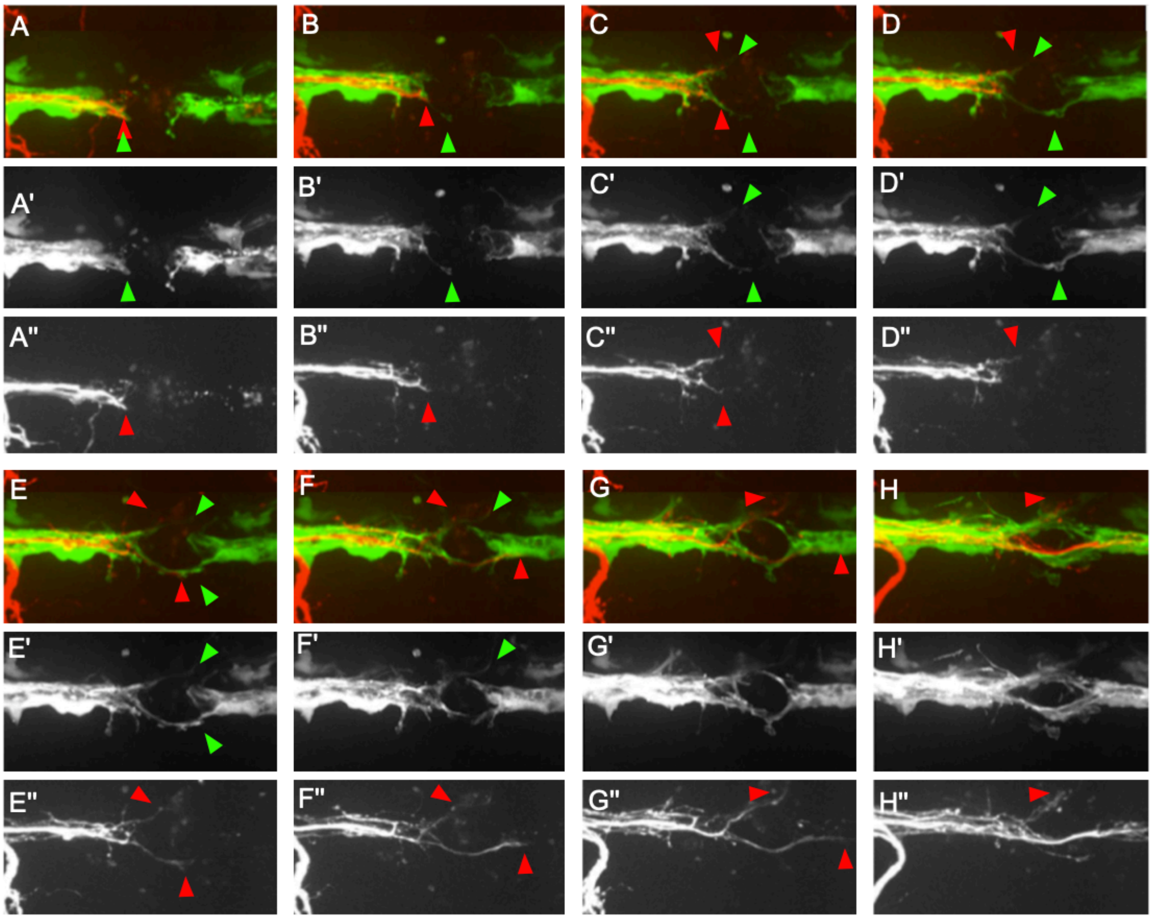


**Supplemental Figure 3.** Series of still images taken from Supplemental Movie 1 (also shown in Figure 6, panel E). They show eight time points during the repair of the gap in the glial scaffold (A'-H') and axonal regeneration after transection (A"-H") in a wild-type animal. Rostral is left and caudal is right. In all panels, the red arrowheads signal the location of the pioneering growth cone of the regenerating axons. The green arrowheads mark the filopodia-like extensions from Schwann cells adjacent to the glial gap. At the start of the series, the axonal terminal stump and the Schwann cells proximal to the gap co-localize (juxtaposition of the green and red arrowhead). In C', a Schwann cell extends a filopodium across the gap, whereas the axons (C") do not grow along this extension of across the gap. In D', several extensions from Schwann cells are clearly visible at the top and bottom aspects of the image. The shape of the lower extension from a Schwann cell did not change shape, suggesting that they are stabilized, perhaps through interactions with the substrate. Proximal axons (D") start to grow along these Schwann-cell protrusions. In E’, the extensions from the anterior and posterior Schwann cells have crossed the gap, physically interact and commence to reconstitute a continuous glial scaffold. The axonal projections (E”), however, have suffered a retraction towards the proximal stump. F’-H’ show a continuation of Schwann cells behavior, increasing the contacts and closing the gap, which is obvious by the smaller area of the gap (distance between proximal and distal Schwann cells). F”-H” show a more robust and persistent extension of the axonal growth cones, which grow nearly strictly along the Schwann cells extensions. Although the gap in the glial scaffold is much reduced, nerve fibers show discrete defasciculation (H”).

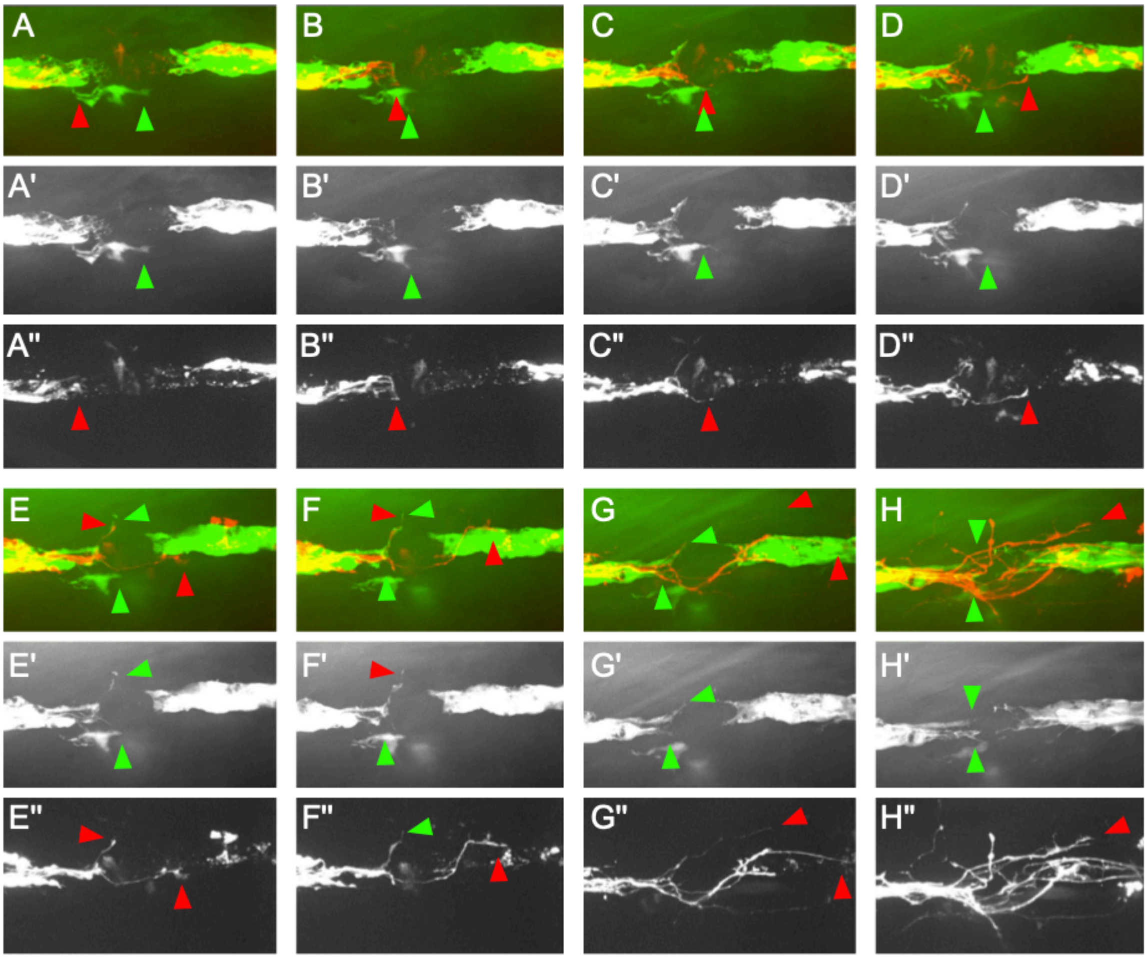


**Supplemental Figure 4.** Series of still images taken from Supplemental Movie 2 (also shown in Figure 6, panel F). They show eight time points during the repair of the gap in the Schwann-cell scaffold (A’-H’) and axonal regeneration after transection (A”-H”), in a Sarm1-mutant specimen. In all panels, the red arrowheads signal the location of the pioneering growth cone of the regenerating axons, and the green arrowheads mark filopodia-like extensions from Schwann cells. Unlike the wild-type situation shown in Supplemental Figure 1, the Schwann cells adjacent to the gap form small filopodia-like extensions, but which never cross the gap. The proximal axon stumps eventually form growth cones that cross the gap at various locations and, upon finding distal Schwann cells, grow along the glial scaffold (E''-H''). Nerve fibers show extensive local defasciculation (H'').
